# Substrate Transport is Mediated not only by P-glycoprotein but also by Lipid Penetration

**DOI:** 10.1101/713503

**Authors:** Yuqi Yu, Jinan Wang, Zhaoqiang Chen, Guimin Wang, Zhijian Xu, Qiang Shao, Jiye Shi, Weiliang Zhu

**Affiliations:** Drug Discovery and Design Center, CAS Key Laboratory of Receptor Research, Shanghai Institute of Materia Medica, Chinese Academy of Sciences, 555 Zuchongzhi Road, Shanghai, 201203, China; University of Chinese Academy of Sciences, UCAS, 19A Yuquan Road, Beijing, 100049, China; UCB Biopharma SPRL, Chemin du Foriest, Braine-l’Alleud, Belgium; Open Studio for Druggability Research of Marine Natural Products, Pilot National Laboratory for Marine Science and Technology, 1 Wenhai Road, Aoshanwei, Jimo, Qingdao, 266237, China

## Abstract

In association with large-scale conformational changes, the members of the ATP-binding cassette (ABC) transporter superfamily such as P-glycoprotein (P-gp) pump endogenous cytotoxic substances and exogenous drugs out of cells. Here, a series of nonequilibrium-driven molecular dynamics (MD) simulations are sophisticatedly combined to provide a generally effective access to quantitatively investigate such a complex biological process that has been posing a great challenge for experiments and computational simulations. Both common features and unique characteristics of multiple ligands (substrates or inhibitors) that are recognized by P-gps from mouse and human species are quantitatively explored, providing additional insights into experimentally suggested ligand transport pathways and summarizing the important roles of not only different P-gps but also lipids in regulating ligand transport. These findings reveal the molecular mechanism underlying the transport of ligands by P-gps from different species and emphasize the consideration of lipid effects on the future design of effective P-gp inhibitors.

## Introduction

P-glycoprotein (P-gp), a representative member of the ATP-binding cassette (ABC) transporter superfamily,^1^ can detoxify cells by pumping a wide range of cytotoxic substances across the membrane.^2^ P-gp is expressed in many “barriers” of the body, including the blood-brain barrier (BBB), kidney, and liver, and plays an important role in the pharmacokinetics and bioavailability of drugs by mediating their transport.^3^ Since the first report in the 1970s that P-gp overexpression in tumor cells increases the expulsion of anticancer drugs, P-gp-induced multidrug resistance (MDR) in many cancers, which is well recognized, has made P-gp a valid therapeutic target.^1,3^ The concurrent administration of anticancer drugs and P-gp inhibitors indicates that efflux modulation with P-gp inhibitors is a promising strategy for tackling MDR problems in chemotherapy.^4,5^ Several of the first-to third-generation P-gp inhibitors, many of which are themselves substrates for ABC transporters that compete with drugs for efflux by P-gp pumps, have been studied in the clinic. Nevertheless, some of these inhibitors have failed due to weak bioactivity and the requirement of using a high dosage with unacceptably high toxicity, but others have shown remarkably better *in vitro* binding affinities and selectivity but were still suspended after phase III clinical trials mainly due to cellular toxicity or reduced efficiency *in vivo*.^5,6^ In addition, differences in the efflux of the same ligand by P-gps from different species represent another serious challenge. An inhibitor with a perfect pharmacological effect in animal models might fail in clinical trials due to its potential to act as a human P-gp substrate. Therefore, a full understanding of the molecular mechanism of substrate/inhibitor recognition by P-gps from different species might facilitate the design of novel and effective P-gp inhibitors to improve the success rate of drug discovery and development.

The structure of P-gp comprises two homologous halves, each consisting of a transmembrane domain and a nucleotide-binding domain. X-ray crystallography experiments of mouse P-gp and several bacterial homologs have revealed two functional states: inward-facing and outward-facing (Figure 1). A popular “ATP-switch model”^7,8^ has been proposed based on the experimentally released structures^9-11^ together with extensive biochemical data^12-18^ to illustrate the general process of P-gp-mediated substrate transport. First, ATPs bind to the two nucleotide-binding domains, leading to the domain dimerization. The transmembrane domains then undergo a conformational change from the inward-facing to outward-facing, which exposes the binding site and pumps the substrate out of the cell. Finally, P-gp restores to the inward-facing conformation after ATPs are hydrolyzed and the relevant products (ADP and Pi) are released. Despite this general description, the important details remain elusive, mainly due to the experimental difficulty in monitoring the global dynamic transition of P-gp. Of particular interest is the mechanism underlying substrate transport inside the structurally dynamic P-gp.

**Figure 1.**
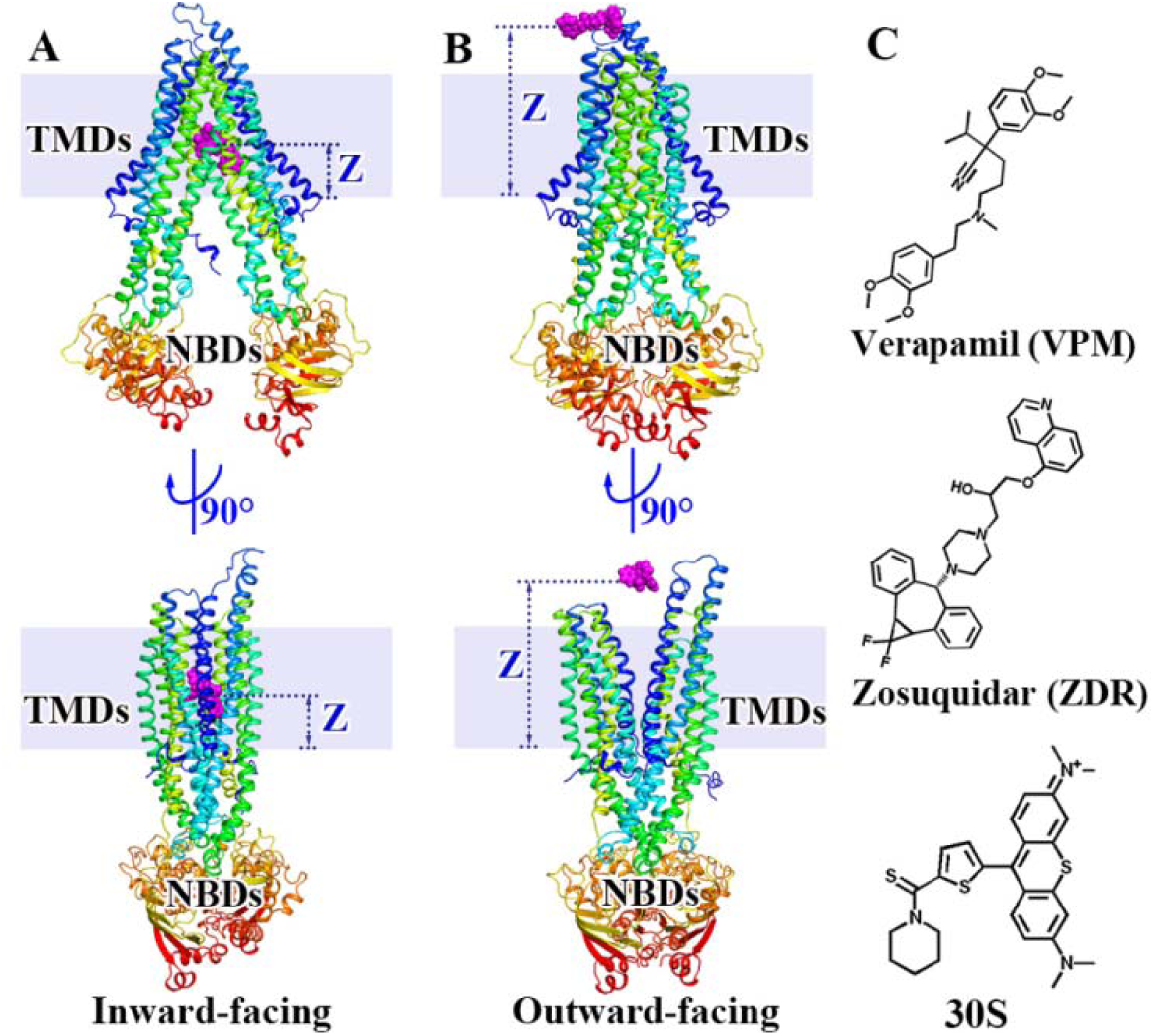
Structures of P-gp and the ligands under study. (A) Inward-facing and (B) outward-facing states of P-gp (TMDs: transmembrane domains; NBDs: nucleotide-binding domains). The ligand positions in parts A and B demonstrate the transport scope from the binding pocket inside P-gp to the extracellular medium. The reaction coordinate Z describing ligand transport is defined as the Z-component distance from the ligand center-of-mass (COM) to the plane of the inner leaflet of lipids (the lipid boundary is shown by a gray rectangle). (C) Structures of verapamil, zosuquidar, and 30S.

Molecular dynamics (MD) simulation represents an alternative approach for investigating questions that are difficult to address directly with experiments. Currently, most simulation research has solely focused on the local structural dynamics of P-gp or the binding interaction analysis of various substrates/inhibitors to P-gp and the lipid bilayer,^19-23^ only a few studies have reported simulated substrate transport events that do not cover the entire translocation process inside the P-gp.^24-28^ Santos and coworkers defined a substrate entrance gate *via* motion pattern analysis^24^ and calculated the potential of mean force (PMF) for substrate translocation from bulk water to the binding pocket of P-gp through the entrance gate.^25^ Subramanian *et al.* used the umbrella sampling technique to calculate the PMFs of morphine and nicardipine molecules moving from aqueous solution to the transmembrane pore without considering P-gp conformational change.^26^ Wise and coworkers used a targeted molecular dynamics (TMD) simulation as well as a docking technique to investigate the drug-and-inhibitor binding interactions in specific intermediate structures of P-gp^27^ and, more recently, to monitor the substrate transport pathways.^28^ Although these studies provide useful information for understanding the efflux mechanism of P-gp, the entire network of transport pathways and related thermodynamic quantities are unavailable.

For the evaluation of a complex biological process in a dynamic protein, a conventional MD or single enhanced sampling MD method might not be sufficiently efficient to explore the entire process within a manageable simulation time. In this study, we attempt to combine multiple enhanced sampling techniques to provide generally effective access for the investigation of substrate transport in association with protein conformational change. As shown in the simulation workflow (Figure S1 in the Supporting Information (SI)), our Normal model analysis-Umbrella sampling Molecular Dynamics (hereinafter referred to as NUMD) combined protocol that was developed previously^29^ is utilized to yield the essential conformational transition pathway of apo-P-gp between the inward-facing and outward-facing states. Subsequent serial nonequilibrium simulations driven with the TMD technique^30^ are performed among P-gp key intermediate states with bound substrate to guide the protein conformational change along the NUMD-generated pathway that mimics a truly dynamic situation for the spontaneous transport of the substrate inside the P-gp. A subsequent steered molecular dynamics (SMD)^31^ simulation ensures that the substrate leaves the P-gp and diffuses into the extracellular medium. Finally, the umbrella sampling approach implemented in NUMD is utilized extensively to calculate the relevant thermodynamic quantities for substrate transport. The selection of key intermediate states according to distinct P-gp transition phases is shown in Figure S2, and a detailed description of the entire workflow and the simulation settings is also provided in the SI Appendix, Materials and Methods.

Comparative studies of different substrates/inhibitors acting in P-gps from different species can enable a comprehensive understanding of how P-gp discriminates substrates and inhibitors.^32,33^ In this regard, three ligands with remarkably different transport properties in mouse and human P-gps are investigated here, among which verapamil (VPM) can stimulate the ATPase activity of P-gp and be transported^34,35^ but zosuquidar (ZDR, also known as LY335979) inhibits ATPase activity at the nanomolar range and can be hardly transported by either type of P-gp,^36,37^ 30S that was discovered through the structural modification of rhodamine can be transported by human P-gp but hardly by mouse P-gp.^38,39^ Thus, VPM can be treated as a substrate but ZDR is an inhibitor of mouse and human P-gps, and 30S is more likely to be an inhibitor of mouse P-gp but a substrate for human P-gp. Systematic studies of the six complex systems (VPM, ZDR, and 30S in mouse and human P-gps) shed light on a set of general principles underlying ligand transport by P-gp and provide valuable information for the development of P-gp inhibitors.

## Results and Discussion

### Connection of Statistically Averaged Microscopic Events to Macroscopic Observations

The distance (Z) from the COM of the ligand to the inner leaflet of the lipid bilayer is used as a suitable reaction coordinate to track ligand transport through P-gp (Figure 1 & Figure S1B). With the combined use of a series of enhanced sampling MD simulations, the motion of all three small molecules (VPM, ZDR, and 30S) through mouse/human P-gps as P-gps undergo conformational changes is mimicked at the atomistic level to provide details of the transport of the ligands from the P-gp binding pocket to the extracellular side. In the TMD simulation part, the ligand trajectories, as represented by the time-series of the Z distance (Figure S3), indicate the spontaneous leaving of the three ligands from the P-gp binding pocket. In the SMD, three independent simulations per system are run to ensure that each ligand completes its translocation to arrive at the extracellular side (Figure S4), and the trajectory with the lowest pulling force (Figure S5) is selected in combination with the TMD trajectories for the PMF calculation.

As shown in Figures 2A-B, the free energy barriers for ZDR transport through P-gp are much higher than those of VPM in both the mouse and human P-gp systems (Δ*G*^*^_ZDR_ ≈ 16.3 kCal/mol and ΔG^*^_VPM_≈ 6.3 kCal/mol in mouse P-gp, ΔG^*^_ZDR_≈ 25.9 kCal/mol and ΔG^*^_VPM_≈ 10.5 kCal/mol in human P-gp), clearly indicating that ZDR is always much more difficult to transport than VPM in P-gps. These results are in strong agreement with experimental observations: ZDR is a much more effective P-gp inhibitor than VPM.^40,41^ Additionally, the lower Δ*G*^*^_VPM_ with mouse P-gp than human P-gp indicates that VPM can be more easily transported by mouse P-gp, which is certainly consistent with the reported larger efflux ratio of VPM (*viz.* can be transported more easily) with mouse P-gp than human P-gp.^42^ Impressively, the difference between P-gps from different species is even clearer for P-gps transporting 30S: The Δ*G*^*^_30S_ with human P-gp (5.9 kCal/mol) is significantly lower than that with the mouse analog (27.2 kCal/mol). Consistently, Detty and coworkers reported that 30S cannot stimulate mouse P-gp ATPase activity but can increase the basal activity of human P-gp by 9-fold (see Tables 1 and S1 in Ref. 38), demonstrating that 30S is much more easily transported by human P-gp. Thus, the present simulations effectively reflect the experimentally reported recognition of substrates/inhibitors by mouse and human P-gps.

**Figure 2.**
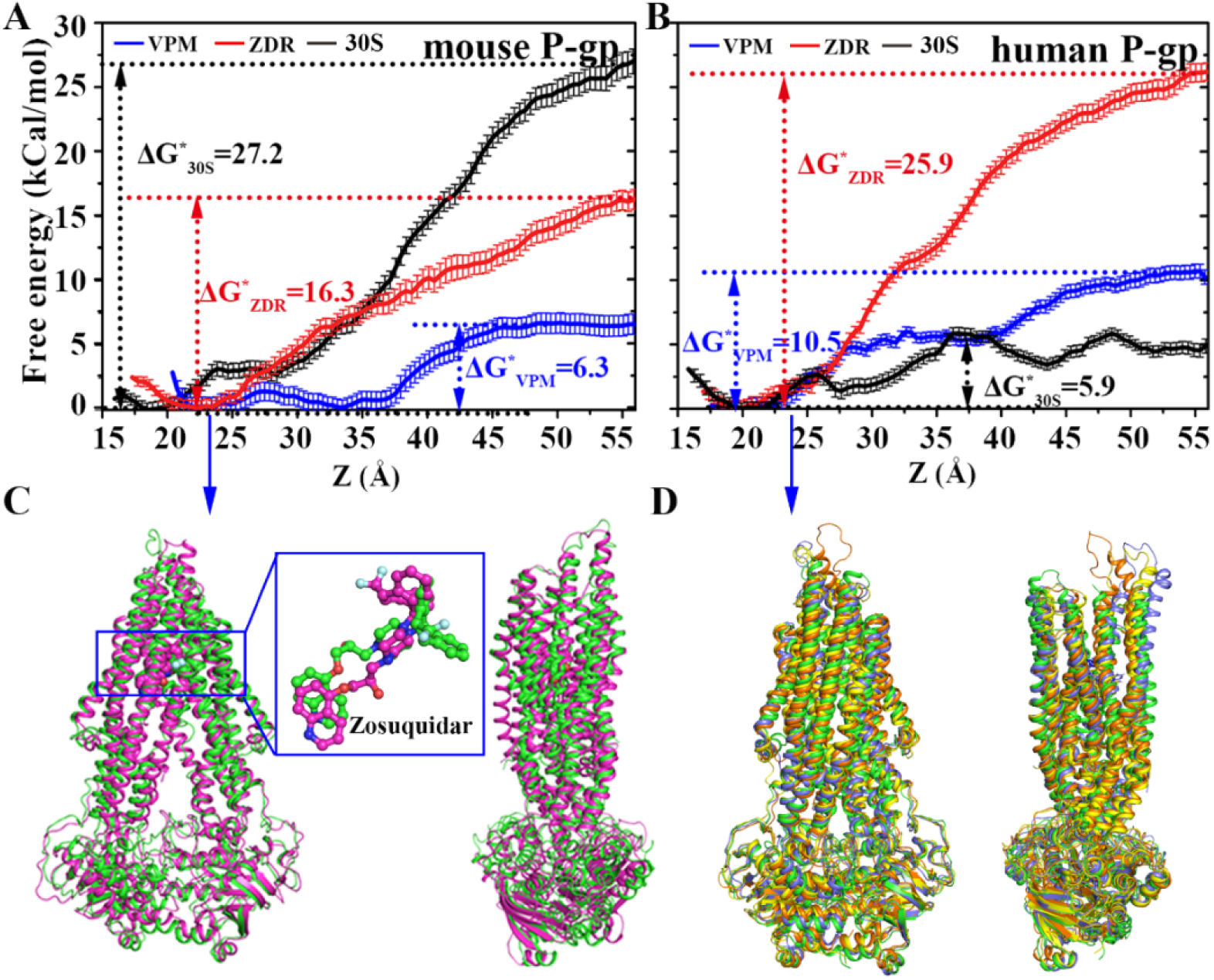
Predicted potential of mean forces (PMFs) for ligand transport and key intermediate structures of P-gp. (A-B) PMFs for ZDR, VPM, and 30S transport through mouse and human P-gps along the Z direction. (C) Superposition of the simulated mouse P-gp-ZDR complex structure at a local minimum of Z ≈ 23.0 Å in panel A (pink) with the cryo-EM experimental structure (PDB code: 6FN1, green). (D) Superposition of the simulated human P-gp structures at a local minimum of Z ≈ 23.0 Å in panel B (VPM/ZDR/30S-bound systems colored in orange, purple, and yellow, respectively) with the cryo-EM experimental apo structure (PDB code: 6C0V, green).

Intriguingly, a cryo-EM structure of the mouse P-gp-ZDR complex was recently reported.^43^ Through structural alignment, we observe that the experimentally determined protein structure is very similar to our simulated structure at the local minimum of Z ≈ 23.0 Å (protein C_α_-atom root-mean-square deviation (RMSD_Cα_) of 2.8 Å) and the ZDR molecule is also captured at a similar position except for the orientation of its heptatomic ring tail (Figure 2C). Similarly, our simulated human P-gp structures near the local minimum of Z ≈ 23.0 Å also resemble the recently released cryo-EM structure^44^ of apo-human P-gp (RMSD_Cα_ = 2.6 ± 0.3 Å), which is characterized by partially opened transmembrane domains (Figure 2D). The consistencies between the experimentally determined intermediate structures and those predicted with the simulations also demonstrate that our strategy and simulation approach are reasonable.

The transport of each ligand passes through a series of residues that characterize the feature of the P-gp transport tunnel. The occupancies of all residues that interact closely with the ligands (≤ 4.0 Å) are calculated (Figures S6 and S7). The key protein residues (occupancy > 50% in either P-gp-ligand complex) and the transport pathway of each ligand are indicated in Figures 3A-B. Twelve and fifteen key residues are found in the mouse and human P-gps, respectively, and 10 of these residues are equivalent (L64↔L65, M68↔M69, F332↔F336, L335↔L339, I336↔I340, F339↔F343, F724↔F728, F728↔F732, F979↔F983, M982↔M986, in mouse or human P-gp), suggesting that the transport of VPM/ZDR/30S by mouse P-gp shares a basic profile with that by human P-gp. Notably, all simulations identified key residues inside human P-pg, except M69, have been reported in the drug translocation pathway investigated by Loo *et al.* using arginine-scanning mutagenesis.^45^ In summary, our simulations achieve strong agreement with experimental results, indicating the high reliability of the present simulations.

**Figure 3.**
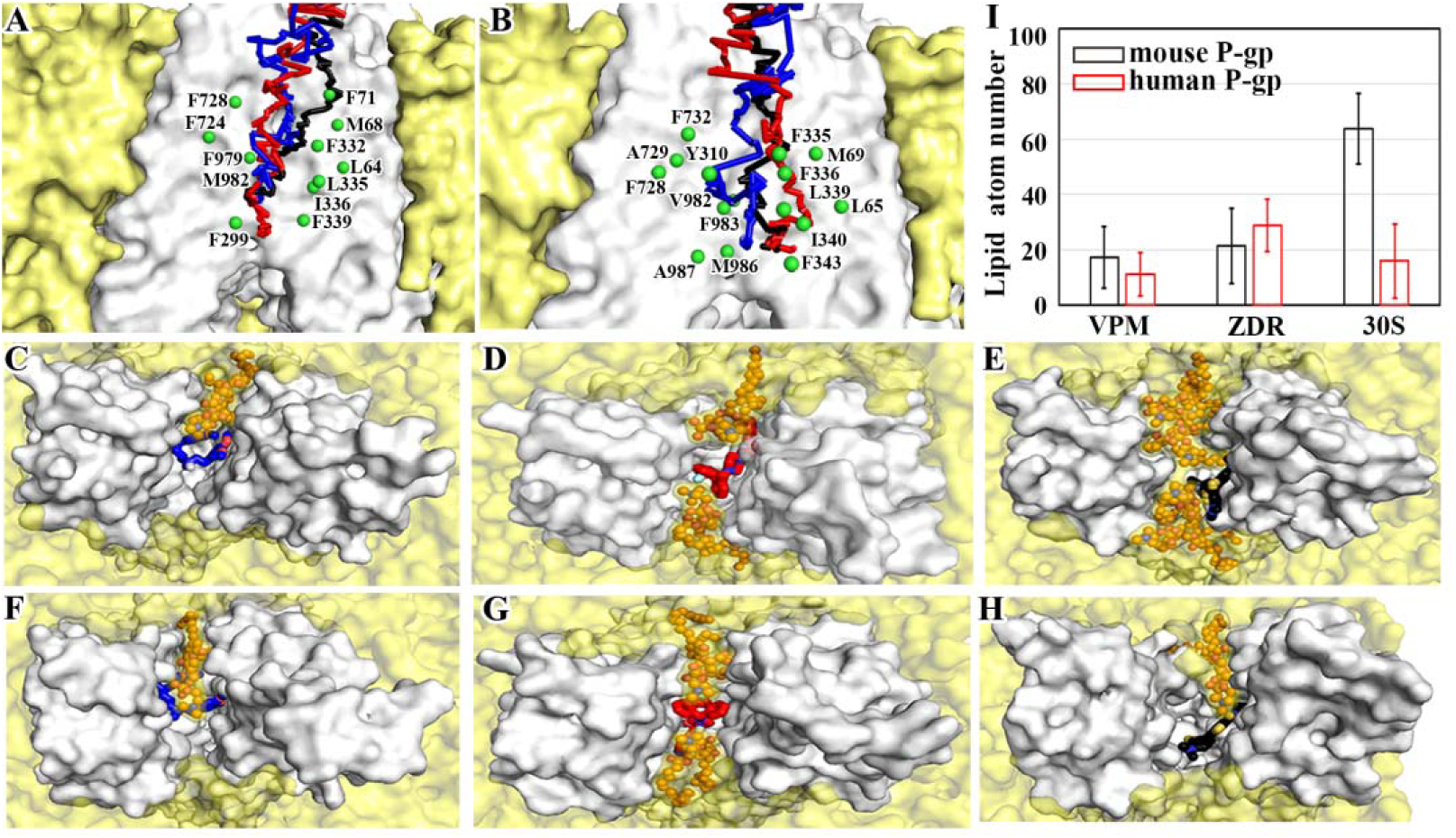
Interaction details between ligands and P-gps during transport. (A-B) Key residues of mouse and human P-gps involved in the transport of VPM, ZDR, and 30S, whose pathways are indicated as blue, red, and black lines, respectively. Snapshot top views indicate the penetration of the lipids into (C-E) mouse and (F-H) human P-gps and their interactions with the ligands. The P-gps and surrounding lipid bilayer are represented by white and yellow surfaces, and the ligands (VPM/ZDR/30S) as well as inserted lipids are shown with blue/red/black and orange sticks. (I) Averaged atom numbers of lipids interacting with the ligands in the respective complex systems.

### P-gp-ligand Hydrophobic Interactions Serve as the Main Driving Force for Ligand Transport

Figure 4 shows the P-gp-ligand interaction map, in which the detailed amino-acid residues interacting with each ligand (≤ 4.0 Å) during its transport process are counted. Hydrophobic interactions (blue/red/black triangles for VPM/ZDR/30S) dominate the P-gp-ligand interactions towards all three ligands, and a small number of hydrogen bonds (green triangles) are also present in the cases of VPM and ZDR. Ligand transport is thus accompanied by the constant formation and breaking of hydrophobic (as well as hydrogen bonding) interactions between the ligand and P-gp. Such interactions have two general tendencies: 1) the total number of ligand-interacting residues decreases during the transport process (inserted panels on the bottom right corner of Figure 4), and 2) the position of Z ≈ 28.0 Å (delimited by the cyan and pink background) seems to be a threshold beyond which the P-gp interactions occur mainly from the position behind the ligand for all complex systems. Therefore, the successive hydrophobic (and hydrogen bonding) interactions formed between each ligand and the P-gp residues work as a driving force that easily draws the ligand to the position of Z ≈ 28.0 Å (corresponding to the low free energies before 28.0 Å in Figures 2A-B). In the late transport stage, the residues ahead of the ligand along the transport pathway cannot allow the formation of sufficient interactions (mainly due to the opening of transmembrane domains) to offset the breaking of the P-gp-ligand interactions formed behind, corresponding to the increasing free energies in Figures 2A-B. The roughly similar residue position dependence of the P-gp-ligand interactions along the transport pathways of all ligands strongly suggests an explanation for the different transport behaviors of the three ligands.

**Figure 4.**
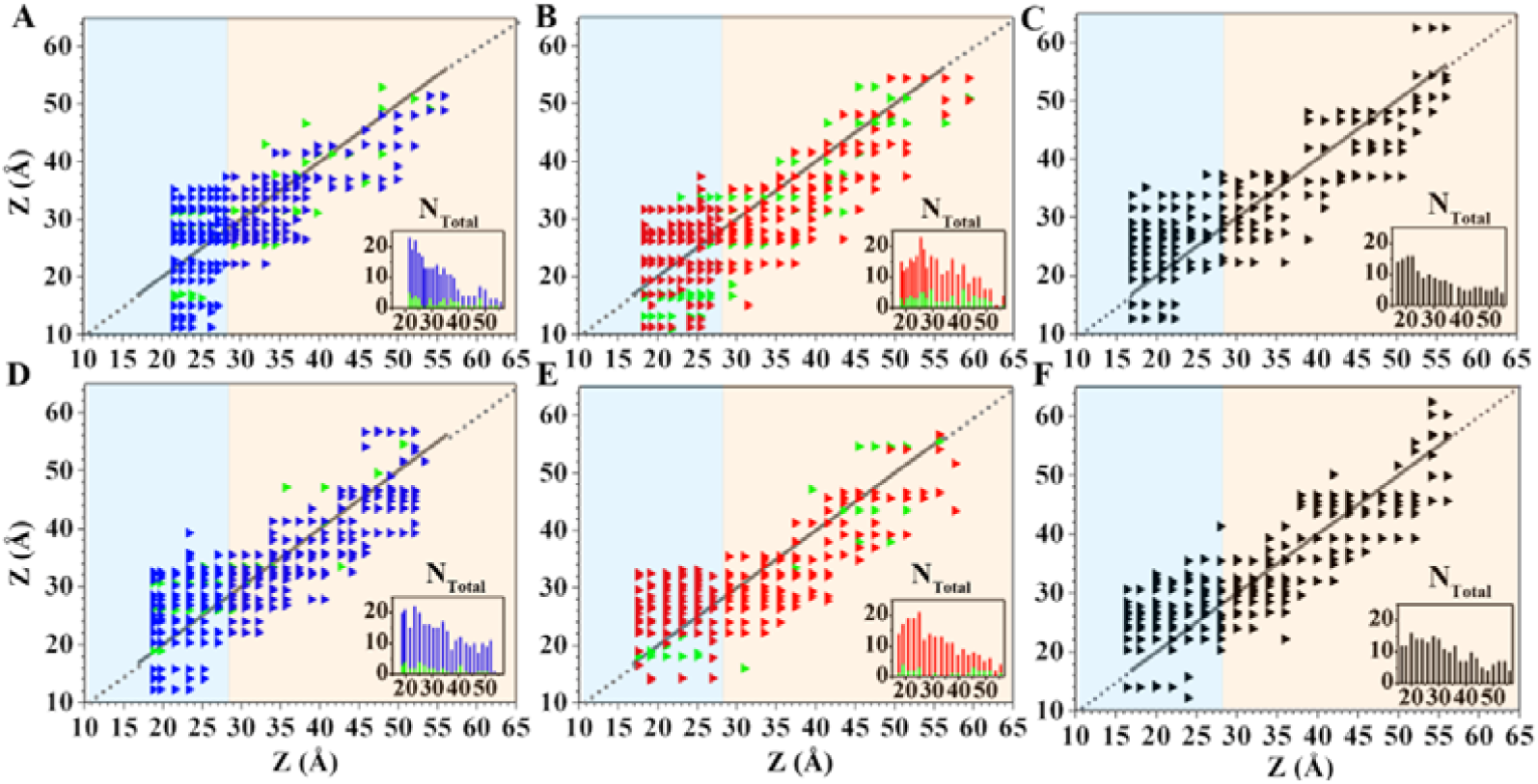
P-gp-ligand interaction maps indicated by the positions of the interacting residues projected in the Z direction (y-axis) *vs.* the ligand COM movement position along the Z direction (x-axis). (A-C) VPM, ZDR, and 30S in mouse P-gp and (D-F) VPM, ZDR, and 30S in human P-gp, respectively. Each triangle represents a residue involved in the ligand interaction. Hydrophobic interactions are indicated with blue, red, or black triangles, and hydrogen bonding interactions are indicated with green triangles. The gray line in each figure denotes the ligand transport pathway along the Z direction. The inserts show the total numbers of ligand-interacting residues along the transport pathways.

### Lipids Penetrate into the P-gp Transmembrane Domains and Interfere with Ligand Transport

A further analysis is performed to explore the ligand-and P-gp system-dependent transport behaviors. A novel phenomenon is generally observed: In the late stage of ligand transport, all P-gp-ligand systems recruit lipid molecules into the central cleft between the P-gp transmembrane domains (see the snapshots in Figures 3C-H and the detailed process in movies S1-S6). Once lipids emerge into P-gps, they interact with the ligands, mainly with the long hydrophobic tails, to hinder their further movement. The average lipid atom numbers interacting with the ligands are provided in Figure 3I. Therefore, the transport of all three ligands in both mouse and human P-gps is affected by the interactions from not only P-gp but also the lipids that penetrate into the P-gp cleft.

To quantitatively compare protein and lipid modulations during ligand transport, the interaction energies between the three ligands to not only P-gp but also emerged lipids are decomposed into van der Waals (vdW) and electrostatic terms, respectively. Apparently, the vdW energy (Figures 5) is predominant over the electrostatic component (Figure S8) in all systems. As shown in Figure 5A, in mouse P-gp, the interactions from the surrounding environment are mainly from the protein at the early transport stage (Z < 28.0 Å). Such P-gp-ligand vdW interactions support ligand transport at this stage, and the interaction strength seems to correlate with the free energy level in Figure 2A (the more negative vdW energies of VPM and ZDR than of 30S corresponding to lower free energies of the former two ligands than the latter before Z = 28.0 Å). Beyond Z = 28.0 Å, the P-gp-ligand vdW interactions are gradually weakened, but lipids also contribute to the interactions toward the ligands. The combined interactions from P-gp and lipids work together to hinder further transport of all ligands. Similar tendencies can also be observed in Figure 5B for the three ligands with human P-gp. In general, in the cases of VPM and ZDR, even though the interaction of P-gp-VPM is slightly more favorable than that of P-gp-ZDR in the case of mouse P-gp but less favorable in the case of human P-gp at the early stage, the combined interactions with P-gp and lipids become less favorable for VPM than for ZDR at the late transport stage, and this finding is obtained for both P-gp systems. 30S, on the other hand, interacts with lipids much more strongly in mouse P-gp than in the human analog. Therefore, the strength of the interactions of the ligands with P-gps as well as lipids correlates well with the accessibility of the ligands to transport, and more favorable vdW interactions with both P-gp and lipids at the late transport stage result in more difficult ligand transport.

**Figure 5.**
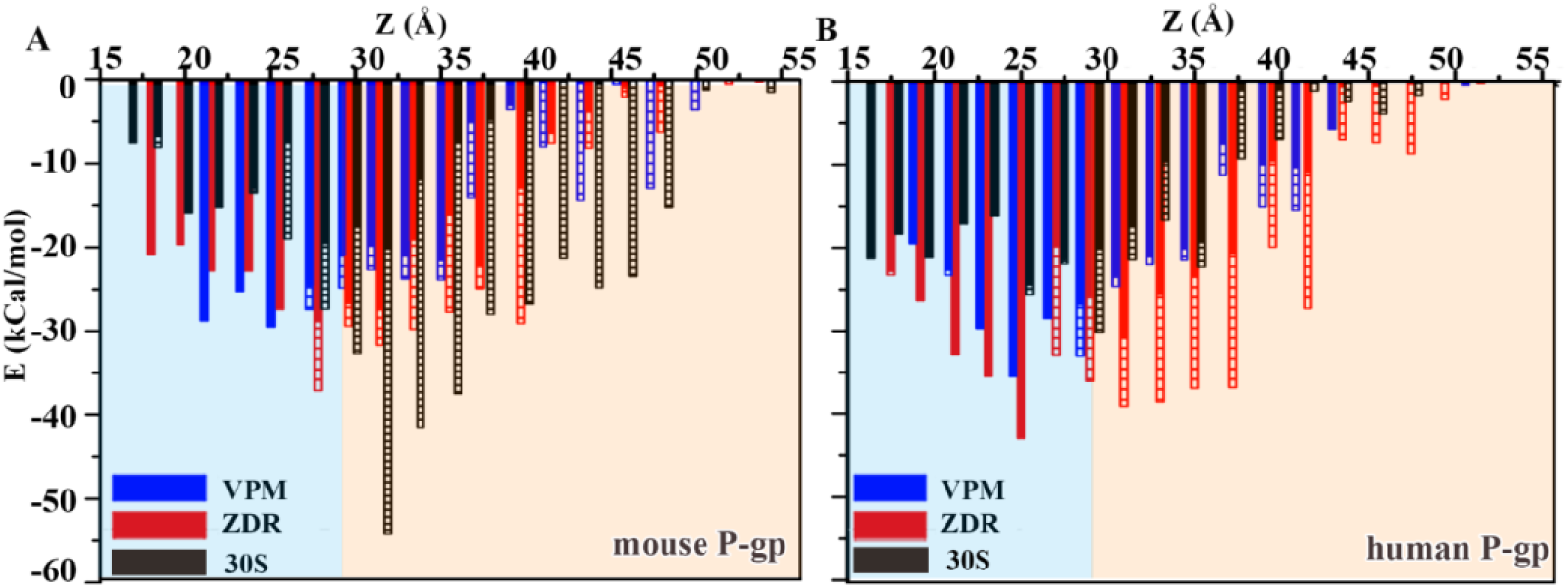
Van der Waals (vdW) energies for the interactions of VPM/ZDR/30S with P-gp residues (solid column) or lipids (dashed column) in (A) mouse and (B) human P-gp systems.

### Hypothesis of Lipid Recruitment according to the Nature of the Ligand and P-gp

While lipid penetration occurs in all complex systems, the greatest differentiation in the level of penetration is obtained for 30S with mouse and human P-gps (Figure 3I). To determine the underlying reason, the transmembrane domain sequences of the mouse and human P-gps are aligned that gives a total of 20 nonconserved residues (Figure S9A). These nonconserved residues are basically distributed in the protein-lipid interface (Figure S9B) and are generally more hydrophobic in mouse P-gp (Table S1), which would supposedly attract more lipids into the transmembrane domain cleft. Additionally, the naturally hydrophobic ligands inside P-gps might also play an important role in recruiting lipids. As reflected by the variance in the nonpolar parts of the solvent-accessible surface areas (SASAs) of the ligand molecules (Figures S10A-B), the three ligands under study display different structural flexibilities during their transport: 30S always maintains its rigid planar structure, which is supposed to be more wide-open to lipids, VPM is extremely flexible, and ZDR is somewhere in between. As a result, the mouse P-gp-30S system recruits the largest amount of lipids during the interaction process. Meanwhile, the presence of P-gp-ligand hydrogen bonding observed with VPM and ZDR but not 30S might, to some extent, restrain the motion of the former two ligands during their transport (Figures S10C-F), which probably impairs the attraction of P-gp and ligand to result in less distinct differentiation in lipid penetration in different P-gps (Figure 3I). Nevertheless, the lipid atom numbers involved in the interactions with VPM are indeed smaller than those with ZDR in both P-gp systems, implying that P-gp inhibitors should be more likely to recruit lipids than substrates.

## Conclusion

Through a systematic investigation of the transport behaviors of three ligands, including substrates and inhibitors, in mouse and human P-gps *via* a combination of a series of nonequilibrium-driven MD simulations, we have been able to more closely connect experiments and simulations. The calculated PMFs for the ligand transport which are capable of quantitatively explaining the experimental observations of the selective transport of VPM but not ZDR by both mouse and human P-gps and 30S transport by human but not mouse P-gp, and the consistency in identifying key P-gp residues along the ligand transport pathway between experiment and simulations support the present simulations.

Based on comparative studies of the six P-gp-ligand complexes that provide similar transport pathways for the test ligands, a model is proposed to demonstrate the transport of ligands inside a P-gp (Figure 6), which is driven by hydrophobic interactions from P-gp. The conversion of the P-gp conformation from the inward-facing to outward-facing state at the late transport stage, however, yields a central cleft between the two transmembrane domains that cannot allow successive P-gp-ligand hydrophobic interactions while leaving room for lipid molecules into P-gp, both obstructing further transport of the ligands. The level of lipid penetration, on the other hand, might depend on the structure and hydrophobic nature of the ligand and the species of P-gp, with a general tendency indicated by the studied cases: fewer lipids emerge into P-gps as the substrates among the test ligands are bound (e.g., VPM in mouse and human P-gps, 30S in human P-gp, Figure 3I). The detailed correlation that may explain the molecular mechanism of substrate/inhibitor recognition by P-gp should be subject to more extensive investigation. Therefore, the difficulty of transporting a ligand is determined by both P-gp and lipid penetration, allowing the easy transport of substrates but making the transport of inhibitors difficult. The future design of effective P-gp inhibitors should take into account the effects of lipid penetration, given the fact that most third-generation inhibitors contain large hydrophobic contact areas.^46-49^ The agreement between the simulated and experimentally determined results also shows that this study provides a feasible approach for predicting whether a ligand is a substrate or inhibitor of a P-gp from a given species.

**Figure 6.**
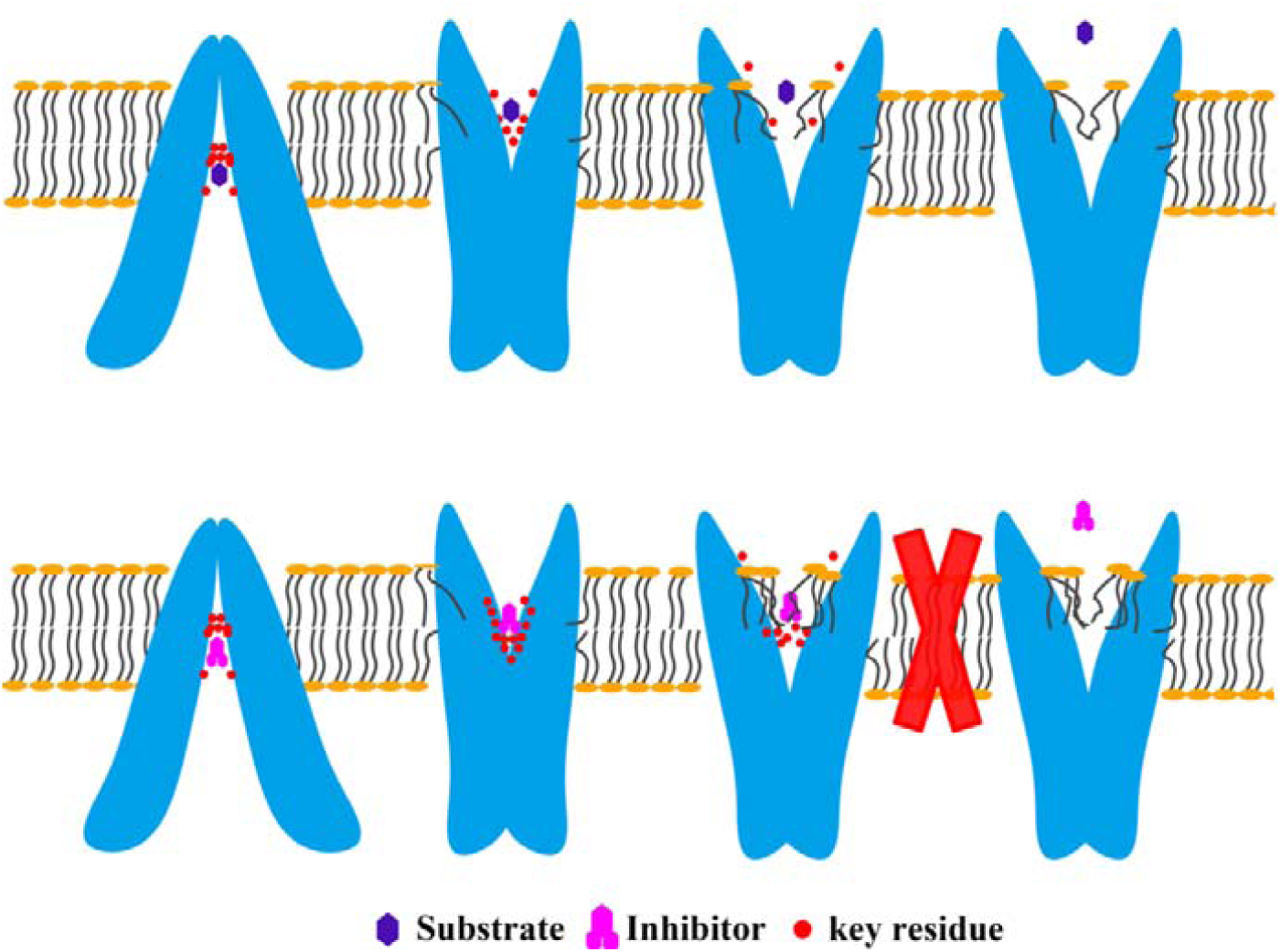
Proposed model for the selective transport of a substrate but not an inhibitor through a P-gp that is determined by the molecular interactions from not only P-gp residues but also lipids penetrated at the late transport stage (generally less obstructive for substrate than for inhibitor).

## Materials and Methods

### P-gp Structure Homology Modeling and P-gp-ligand Complex System Preparation

The inward-facing and outward-facing structures of mouse and human P-gps were modeled using Modeller^50^ based on the crystal structures 4M1M^51^ and 2ONJ,^52^ respectively. For the P-gp-ligand complex systems, the ligands (VPM/ZDR/30S) and ATP were parameterized using a generalized Amber force field (GAFF).^53^ The protein, palmitoyloleoylphosphatidylcholine (POPC) bilayer, and ions were modeled using CHARMM36 force field,^54^ and water was based on the TIP3P model.^55^

### Nonequilibrium MD Simulations Yielding Ligand Transport Pathways

The conformational transition pathway of mouse or human P-gp from its inward-facing state to outward-facing state was generated by NUMD,^29^ and during this simulation, multiple key intermediate structures covering the important conformational transition phases of P-gp were specifically selected as reference structures for the subsequent TMD simulations.^30^ For each type of P-gp, the P-gp-ligand complex system was constructed in an inward-facing state that underwent minimization, heating, and short-time equilibrium (at 310 K) processes with conventional MD using GROMACS version 5.1.1.^56^ Continuous TMD simulations were then run using GROMACS 5.1.1-PLUMED 2.1.2,^57^ with the biased force applied on the protein only to guide the P-gp conformational change among the abovementioned selected intermediate structures that mimic a dynamic situation for the spontaneous transport of ligand inside P-gp. To ensure that each ligand eventually moved away from the P-gp, an additional SMD simulation^31^ was performed on the ligand in the last TMD simulation frame to simulate the diffusion process of the ligand into extracellular medium.

### Ligand Transport PMF Calculation

The calculation of the ligand transport PMF was performed using the umbrella sampling approach embedded in NUMD. A total of ∼100 windows, with a 0.3-0.5 Å interval in the Z distance between neighbor windows, were used for each P-gp-ligand system. The weighted histogram analysis method (WHAM)^58^ was subsequently utilized to calculate the unbiased distribution from the biased sampling and compute the PMF along the reaction coordinate of the Z distance. A detailed description of all abovementioned simulation steps is provided in SI Appendix, Materials and Methods.

## Supporting information

SupportingInformation_text

## Supplementary Information

The supporting information includes the detailed description of Materials and Methods, Figures S1-S13, Tables S1-S2, and videos showing the ligand-lipid interactions during the transport process.

## Acknowledgments

This work was supported by the National Key Research and Development Program (Grant 2016YFA0502301), National Basic Research Program (Grant No. 2014CB910400), and National Natural Science Foundation of China (Grant No. 81573350). This project was partially a SIMM-Pfizer joint project titled “Molecular recognition of ABC family transporters and computational modeling of P-gp transporting mechanism” started on September 2010, and we thank Dr. Xinjun Hou for stimulating discussions. The simulations were run with the TianHe-2 supercomputer in Guangzhou, supported by the Special Program for Applied Research on Super Computation of the NSFC-Guangdong Joint Fund (the second phase) under Grant No. U1501501.

---

**TOC Graphic**

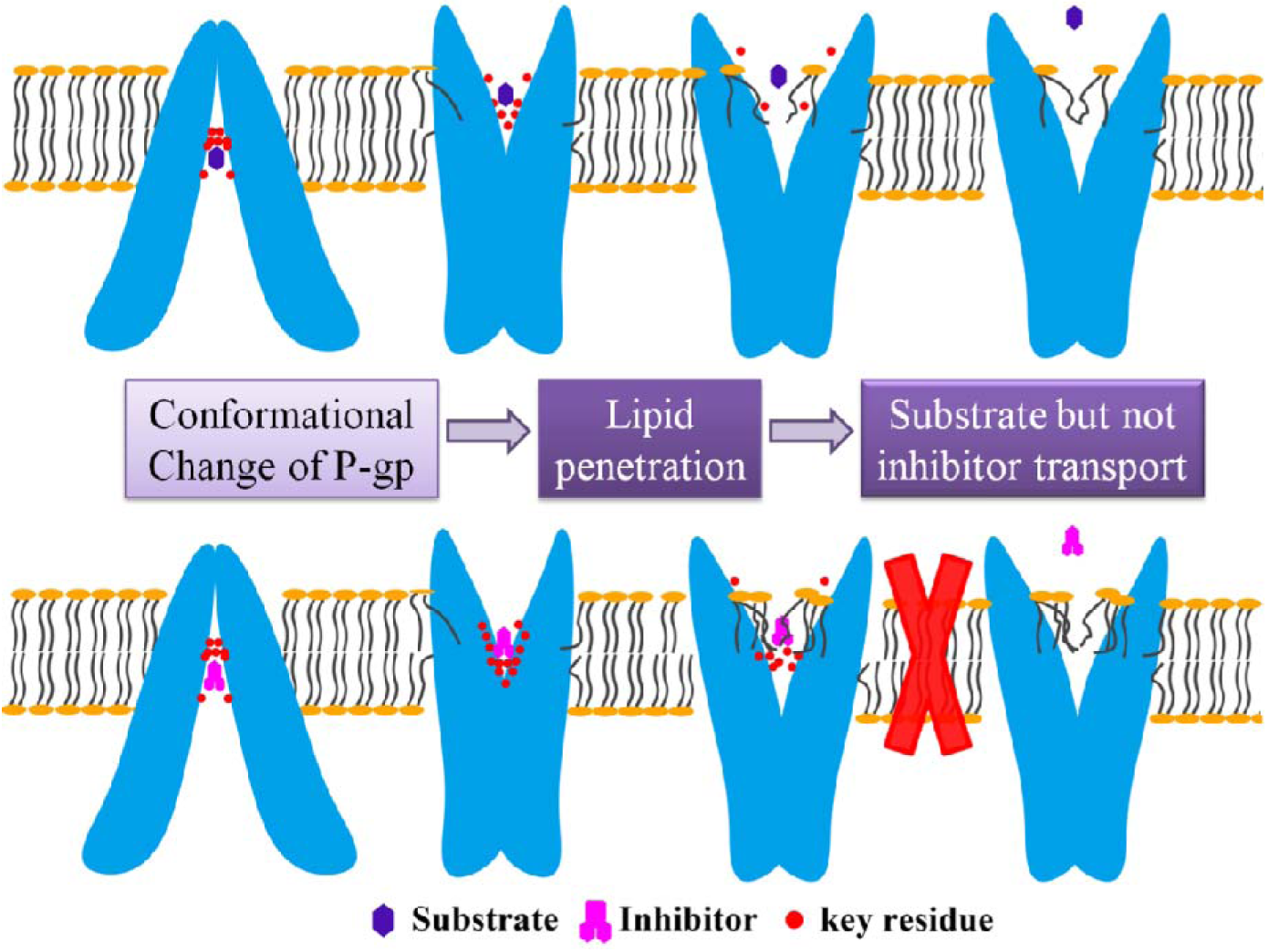

