## SupportingInformation_text for "Substrate Transport is Mediated not only by P-glycoprotein but also by Lipid Penetration"

**This file includes:**

Supporting information text

References for SI reference citations

Figures S1 to S13

Tables S1 to S2

Captions for movies S1 to S6

**Other supplementary materials for this manuscript including the following**:

Movies S1 to S6

**Supplementary Information Text**

**Materials and Methods**

**Homology Modeling for P-gp Structures.** For mouse P-gp, its inward-facing structure is available from the crystal coordinates (PDB code: 4M1M)^1^ and outward-facing structure has been modeled by Stephen G. Aller group based on the crystal outward-facing structure of a bacterial ABC transporter (*S. aureus* Sav1866).^2^ Meanwhile, the inward-facing and outward-facing structures of human P-gp were modeled based on the corresponding structures of mouse analog. Sequence alignment and homology modeling were done by using Modeller.^3^ Sequence identity was 88.7% after the removal of residues 1-30 at N-terminus, 1275-1280 at C-terminus, and linker residues 631-684 that are not available in template sequence (Figure S11). The yielded inward-facing and outward-facing models of human P-gp are quite similar to the corresponding templates of mouse P-gp (RMSD_Cα_ < 0.4 Å). The side-chain of the two structures were then minimized using Maestro^4^ and then evaluated by MolProbity server^5^ (Figure S12), showing high qualities of the built models.

**Conformational Transition Pathways of apo-P-gps Yielded by NUMD and the Selection of Key Intermediate Conformations.** The detailed theoretical description of NUMD method has been given in our previous literature,^6^ and its reliability in measuring protein functional conformational transition pathway and calculating the relevant thermodynamic quantities has been testified in multiple protein systems.^6^^,^^7^ Using the inward-facing structure that is obtained from either X-ray crystallography experiment or homology modeling as a starting endpoint, multiple NUMD iterations were run until the protein arrived at the target outward-facing structure.

In the *k^th^* iteration, the P-gp intermediate structure
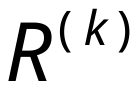
 is generated by:

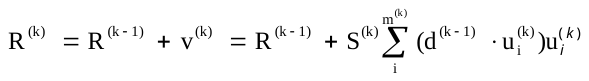
 (1)

where
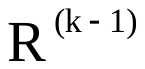
 is the intermediate structure obtained in the (*k-1)^th^* iteration,
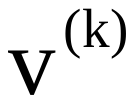
 is the combined displacement along
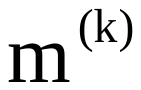
 low-frequency eigenmodes calculated by the normal mode analysis approach implemented in NUMD.^6^ The displacement along the *i^th^* eigenmode is proportional to the projection
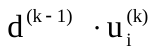
 of the instantaneous distance vector
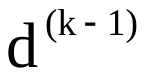
 on
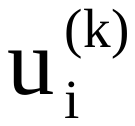
, and scaled by the step size
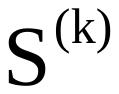
 (10.0). The number of
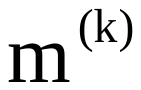
 at *k^th^* iteration is determined by the following function:

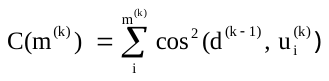
 (2)

The value of
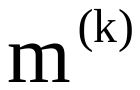
 is the minimal number of modes which start from the lowest frequency mode and end until
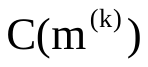
 reaches the cutoff of 0.8 as defined in multiple pioneering studies.^8^^,^^9^ Every intermediate structure obtained in each iteration of NUMD was subject to 1000 minimization steps by conventional MD, and the optimized structure was then used in the next NUMD iteration.

Multiple representative intermediate structures of P-gp were then chosen from all NUMD yielded ones to work as the reference structures in the subsequent non-equilibrium MD simulations to guide the P-gp conformational change. Several reaction coordinates were defined to monitor the global conformational change of P-gp: 1) the COM distance between the two nucleotide-binding domains (NBD1-NBD2) to measure their relative motions, 2) the inter-residue distances of 79/80-736/686, 325/329-968/972, 207/211-850/854 to show the movement of the inner gate, middle gate, and outer gate in the transmembrane domains, respectively, as defined by Pan and Aller^2^ (Figure S13). As shown in Figure S2, the conformational change of P-gp follows step-by-step sequence, in which the abovementioned structural elements move in a non-concerted manner. 4 or 3 key intermediate structures where inflection point occurs along either reaction coordinate were chosen for mouse or human P-gp eventually to cover the entire P-gp conformational change (Figure S2).

**Protein-ligand Complex System Preparation for Non-equilibrium MD Simulations.** It is noteworthy that the abovementioned NUMD yielded intermediate structures work as the reference structures only and are not involved in the preparation of P-gp-ligand complex system. As matter of fact, only the inward-facing structure of P-gp was used as the initial structure for the construction of P-gp-ligand complex system for the non-equilibrium MD simulations. For each P-gp-ligand complex system (VPM, ZDR, or 30S in mouse or human P-gp), the ligand was docked into the ligand-binding pocket of the inward-facing structure of P-gp using Schrödinger software. Two ATPs were also docked into the ATP-binding pockets of NBDs. Protein was prepared using protein preparation wizard (PrepWizard) in Maestro.^4^ Three-dimensional structures of VPM and ZDR were downloaded from PubChem (<https://www.ncbi.nlm.nih.gov/pccompound>), and 30S was drawn and minimized in Maestro.^4^ The three molecules were prepared using Ligprep wizard^10^ and their protonation states at PH 7.0 ± 2.0 were determined using Epik.^11^ The initial docking box center coordinate was settled by referring center of mass for ligand in murine P-gp complex crystal structure (PDB code: 3G60^12^). Molecular docking was performed using Glide module with XP mode.^13^ Poses with lowest binding affinities were selected.

Next, each docked P-gp-ligand complex system (mouse P-gp-VPM, mouse P-gp-ZDR, mouse P-gp-30S, human P-gp-VPM, human P-gp-ZDR, or human P-gp-30S) was inserted into a 125 Å×110 Å palmitoyloleoylphosphatidylcholine (POPC) bilayer. Lipids located within 1 Å of the protein were manually removed. The system was then solvated by TIP3P waters^14^ with 0.15 M NaCl. CHARMM36 force field^15^ was used to model protein, membrane, and ions. The ligand and ATP were parameterized using generalized Amber force field (GAFF)^16^ with restrained electrostatic potential (RESP)^17^ partial charges calculated by fitting to their electrostatic potential from Gaussion 09.^18^ The detailed composition of each system is summarized in Table S1.

**Ligand Transport Pathway Measured by Non-equilibrium MD Simulations.** Each constructed P-gp-ligand complex system was subjected to the preprocesses including the minimization, heating-up, and short-time equilibrium at 310 K with conventional MD performed by GROMACS version 5.1.1.^19^ Firstly, membrane-embedded protein inward-facing structure was minimized to remove steric clash. Then 5 ns MD simulation was run to heat the system from 0 K to 310 K with constraint applied to keep solute fixed. The whole system was then relaxed step by step in NPT ensemble as follows: 500 ps for relaxing solvent, 500 ps for lipid tails, 10 ns for lipid heads, 4 ns for protein side-chain, 10 ns for protein main chain, and 2 ns for ligands and ATP, respectively, giving a total of 27 ns to equilibrate each simulation system. In these processes, temperature was maintained at 310 K using Nosé-Hoover scheme^20^^,^^21^ with coupling time of 1.0 ps. Pressure was maintained at 1 bar using Berendsen algorithm^22^ coupled every 5.0 ps and compressibility was set as 4.5 × 10^-5^ bar^-1^. Bonds to hydrogen atoms were constrained using the LINCS^23^ method every 2 fs. The cutoff for Lennard-Jones interactions was set to 12 Å and the electrostatic interactions were calculated using the particle mesh Ewald (PME) algorithm^24^ with a real-space cutoff of 12 Å.

Targeted molecular dynamics (TMD) simulations^25^ were run for each equilibrated system using GROMACS 5.1.1-PLUMED 2.1.2^26^ to guide the complete conformational change of P-gp along the reasonable pathway starting from the inward-facing to the end of outward-facing that should provide access to the spontaneous transport of ligand. To do so, multiple iterations of TMD simulations each lasting ~20 ns (5 TMD simulations for mouse P-gp or 4 for human P-gp) were performed one after another among the references of predefined P-gp intermediate structures (Figure S2). In each TMD simulation iteration, the P-gp protein was pushed towards its target conformation (selected from NUMD simulation snapshots) by an artificially added restraint potential on all protein C_α_ atoms (*k* = 48.0 ~ 11950.0 (kCal/mol)/A^2^) whereas the ligand (VPM/ZDR/30S) moves freely. Once the target conformation is arrived, the last simulation frame is then used as the initial structure for the next TMD simulation iteration to reach the desired target conformation.

In the case that a ligand cannot by itself transports to the extracellular side at the end of the sequential TMD simulations, additional steered molecular dynamics (SMD) simulations^27^ were run on the last TMD simulation frames to stimulate the diffusion of the ligand into the extracellular medium. To minimize the influence of the artificially added force in SMD on ligand translocation, for each simulation system, 3 frames at the end of the TMD simulation were chosen as the initial structures for independent SMD simulations, with varied atom coordinates and velocities while the configurations of protein and ligand remaining little changed. A very slow velocity of 0.01 Å/ns and a weak biased force with the force constant of *k* = 2.0 (kCal/mol)/ Å^2^ was added on Z-distance to achieve better equilibration in SMD. The trajectory with the smallest pulling force (meaning the easiest to diffuse into the extracellular medium) was chosen for next step calculation.

**Potential of Mean Force (PMF) Calculation for Ligand Transport.** The calculation of the PMF of the ligand transport through P-gp was made using umbrella sampling embedded in NUMD. A total of ~100 snapshots were selected from the abovementioned TMD and SMD simulations for each system with a 0.3-0.5 Å interval of Z distance for the ligand between neighboring snapshots. The windows in the umbrella sampling thus can cover the entire ligand transport pathway. In each window, the P-gp-ligand complex conformation was fist minimized and equilibrated at NPT ensemble for 1 ns, with restraints adding to the ligand and protein C_α_ atoms. Then umbrella sampling was performed for 15 ns per window with a biasing harmonic potential adding to the reaction coordinate of Z distance. Force constants were set as 2.0 ~ 9.0 (kCal/mol)/A^2^ depending on the detailed systems. Weighted histogram analysis method (WHAM)^28^ was finally utilized to calculate the unbiased distribution from the biased sampling and compute the PMF along the reaction coordinate of Z distance.

**Residue Occupancy Calculation.** The occupancy of individual P-gp amino acid residues to interact with the ligand was calculated to figure out the key residues for ligand transport. During the ligand transport from the P-pg binding site to the extracellular medium, a residue was considered as interacting with the ligand as the minimal distance of heavy atoms between the residue and ligand is less than 4 Å. The occupancy of residue *i* was then defined as:

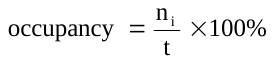
 (3)

where *n_i_* is the number of snapshots in which the residue *i* is within 4 Å around the ligand and *t* is the total number of analyzed snapshots.

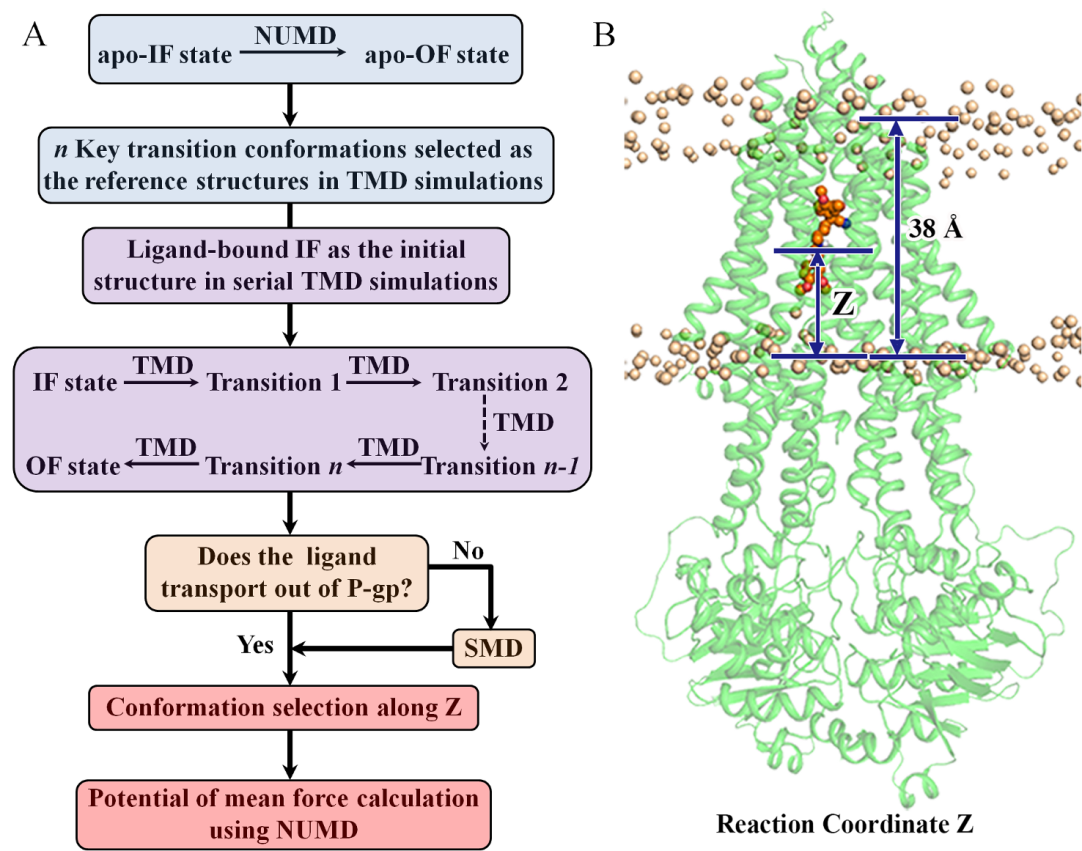

**Figure S1**. (A) Simulation workflow. IF: inward-facing state; OF: outward-facing state; NUMD: normal mode analysis-umbrella sampling molecular dynamics; TMD: targeted molecular dynamics; SMD: steered molecular dynamics. (B) Schematic diagram for the definition of reaction coordinate Z for ligand transport: Z-component distance from ligand center-of-mass (COM) to the plane of inner leaflet lipids. 38 Å represents the total length of the lipid bilayer.

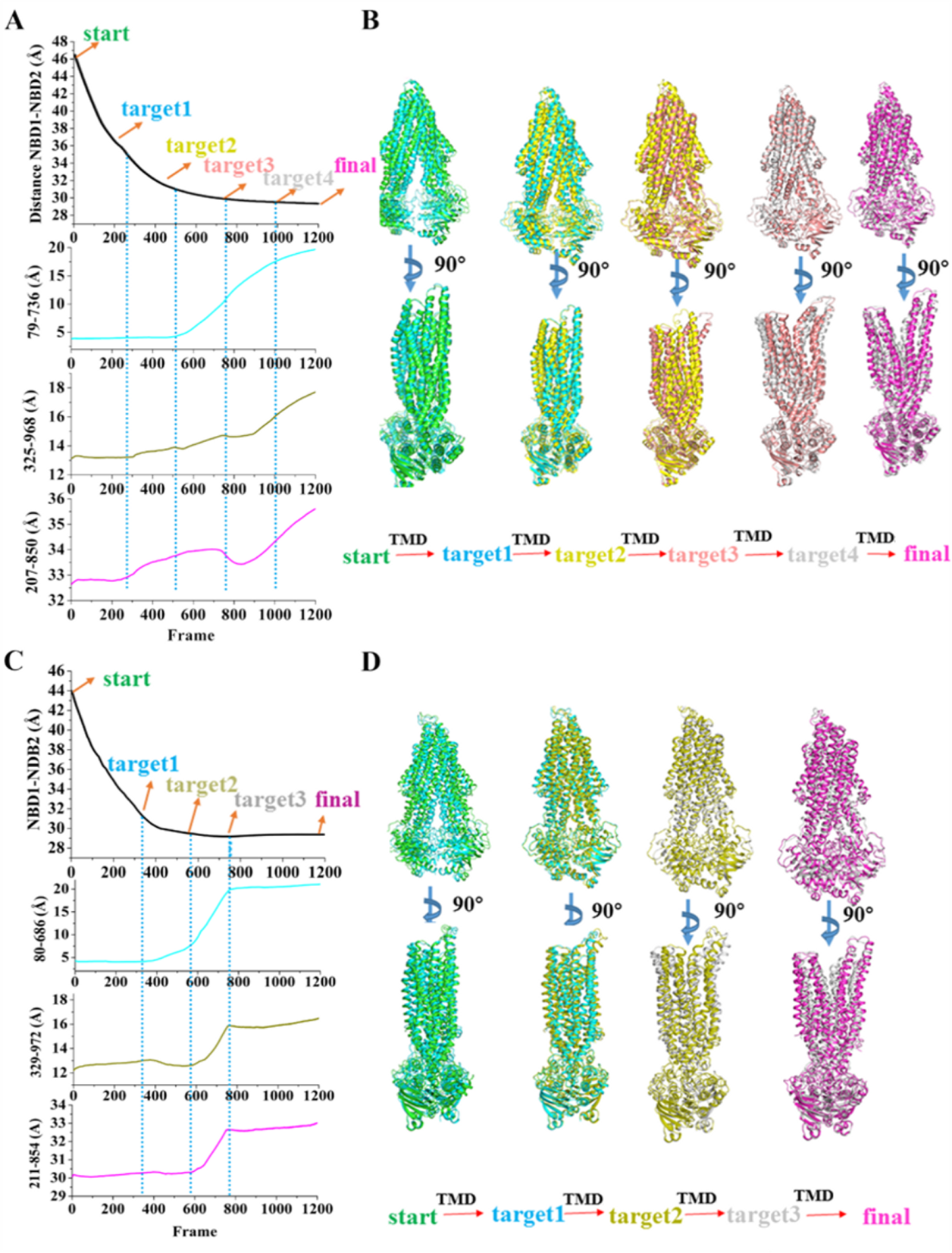

**Figure S2**. The distance changes of P-gp dimeric structural segments (NBD, inner gate, middle gate, and outer gate, see the definition in Figure S13) along the NUMD yielded conformational transition pathways to indicate the distinct transition phases, and the key transition conformations chosen from all transition phases for the subsequent target molecular dynamics (TMD) simulations for (A-B) mouse and (C-D) human P-gp systems, respectively.

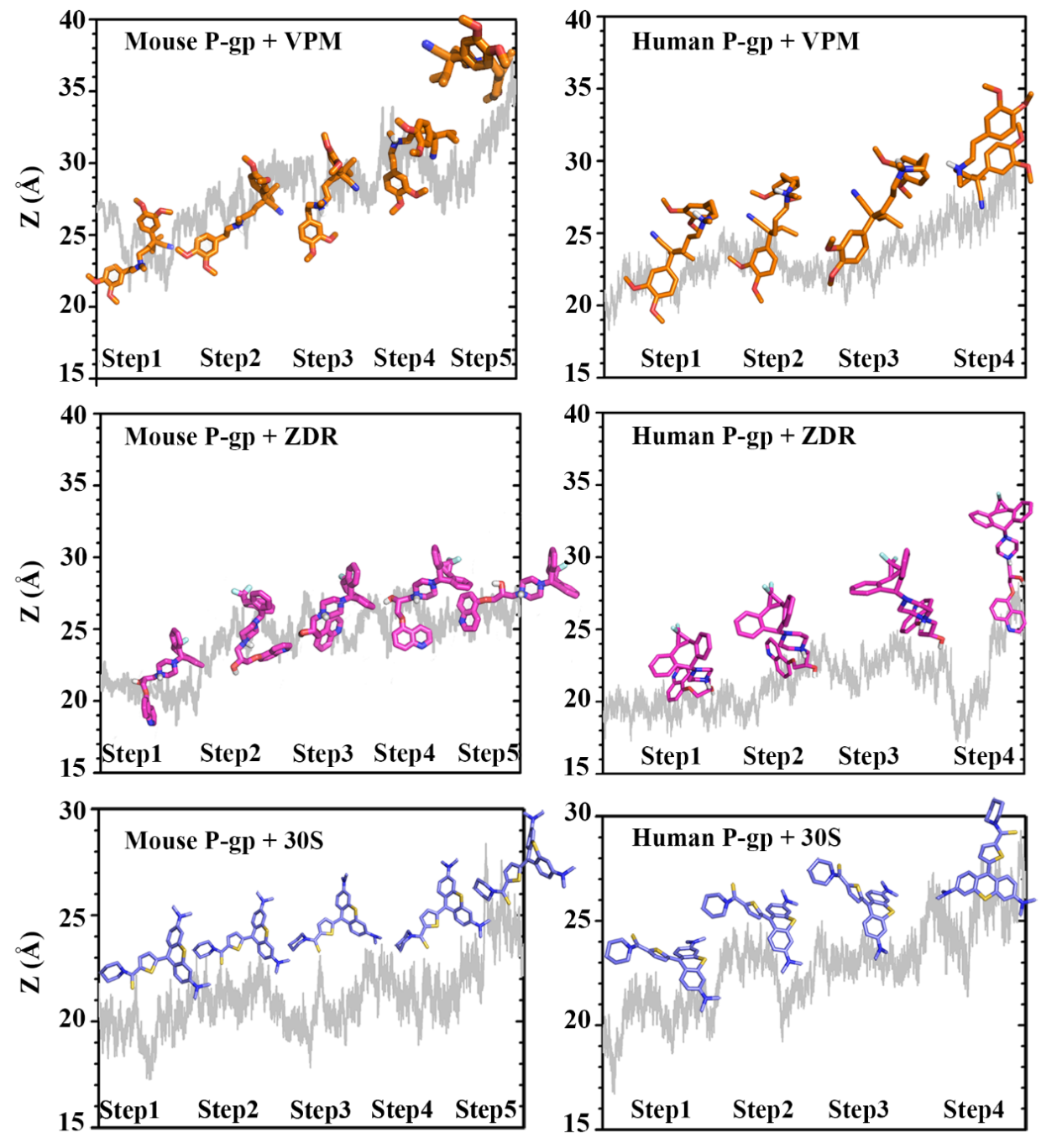

**Figure S3**. Z-component distance for verapamil (VPM), zosuquidar (ZDR), and 30S in mouse and human P-gp systems in multiple steps of TMD simulations. The initial posisiton of the ligands is in the binding packet of P-gp transmembrane domain where the ligands are placed by docking technique.

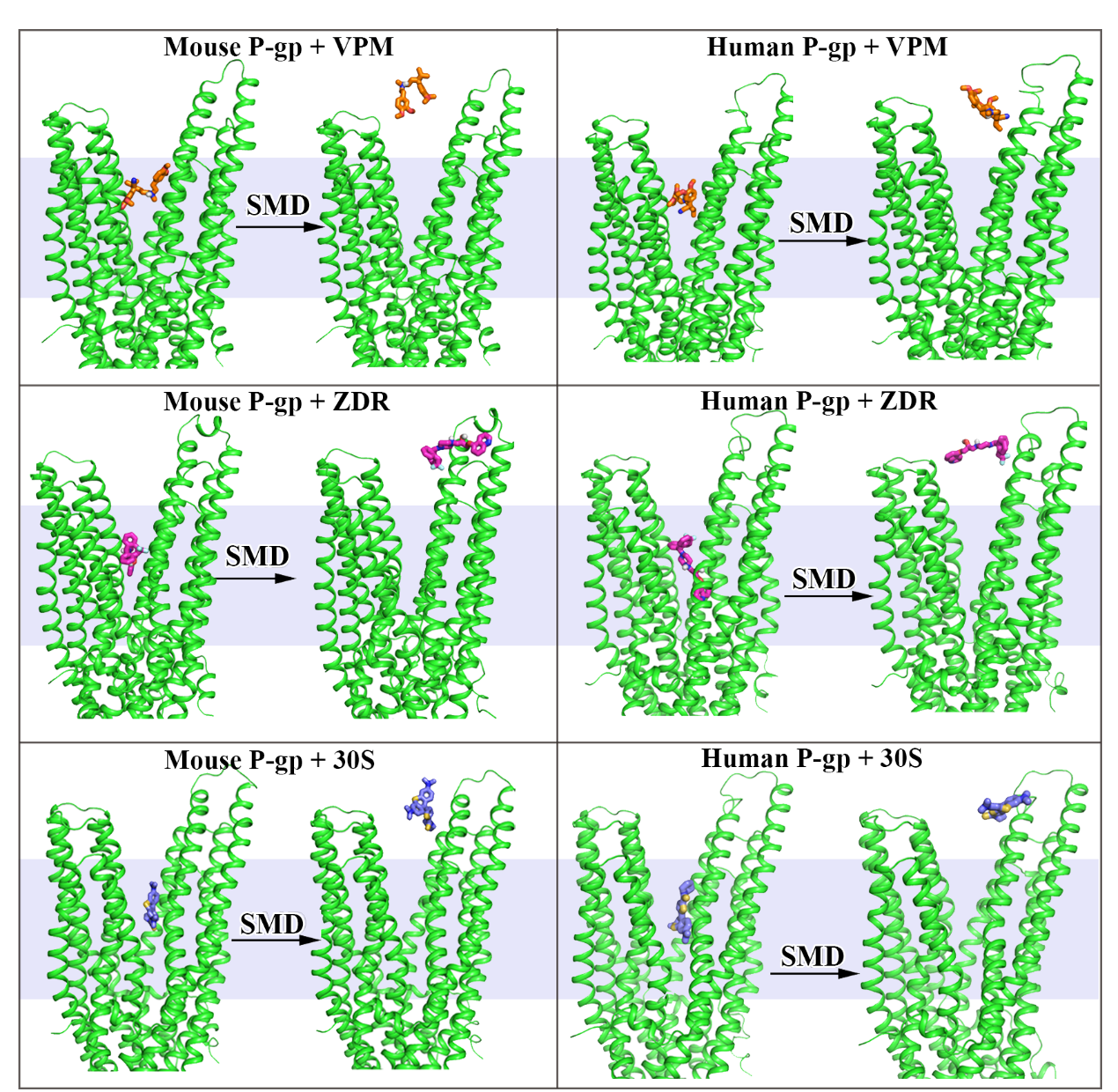

**Figure S4**. The starting and ending ligand positions in the SMD simulations of verapamil (VPM), zosuquidar (ZDR), and 30S in mouse and human P-gp systems, respectively.

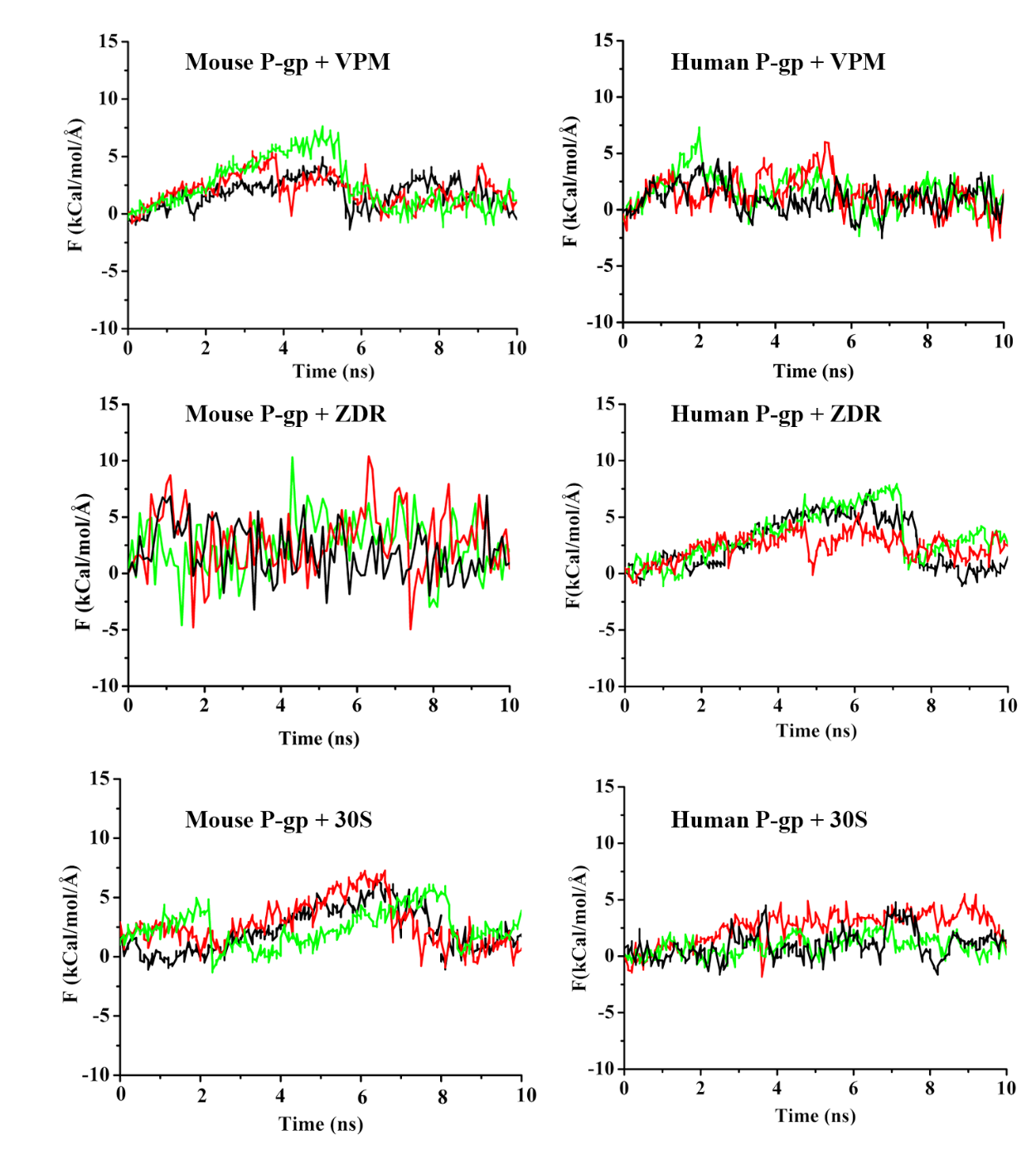

**Figure S5**. Pulling forces of three independent SMD simulation trajectories for verapamil (VPM), zosuquidar (ZDR), and 30S in mouse and human P-gp systems. The trajectory with lowest peak of pulling force in each system (black line) is used for umberlla sampling calculation.

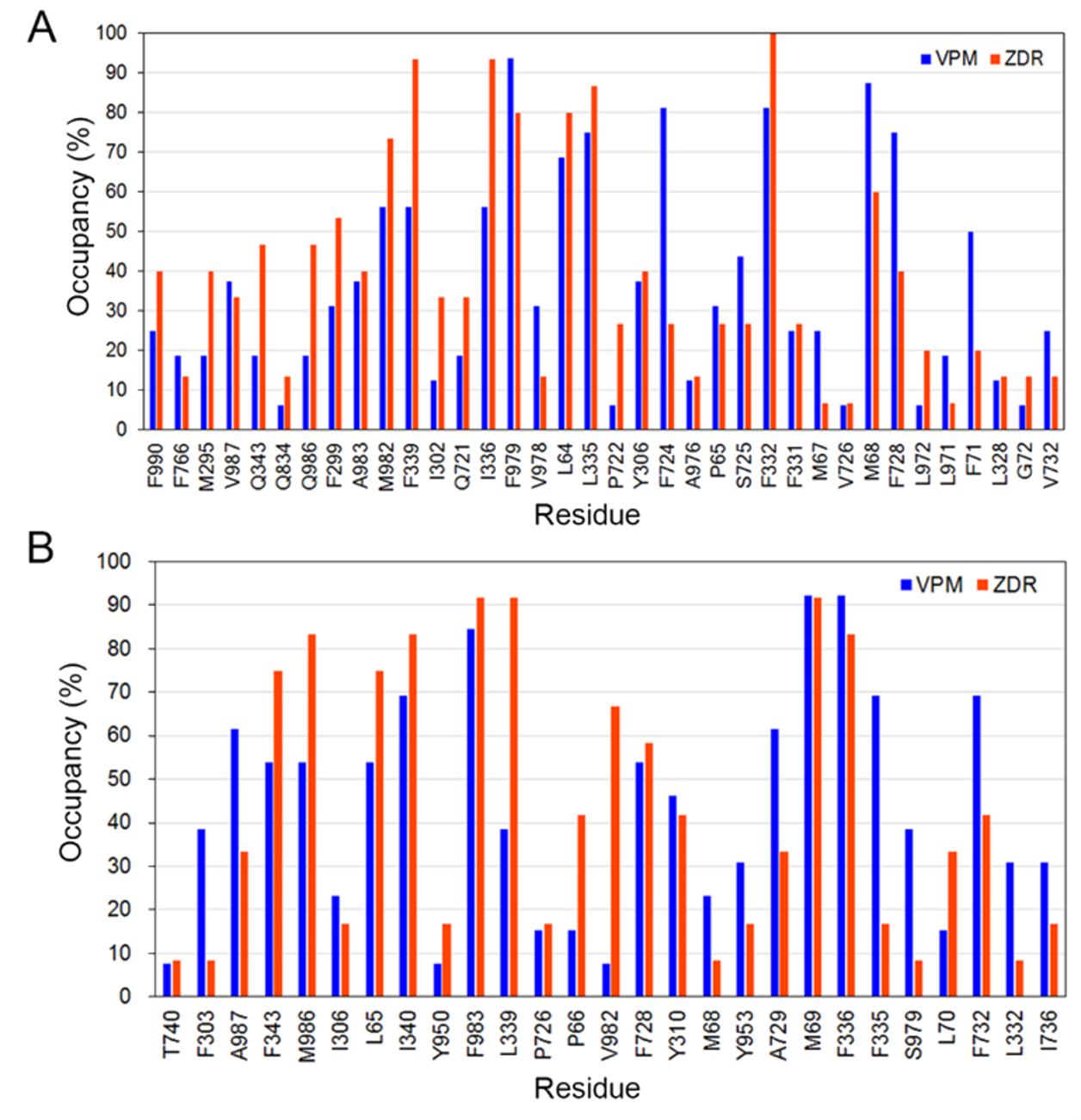

**Figure S6**. Residue occupancy for VPM/ZDR transport in (A) mouse and (B) human P-gps.

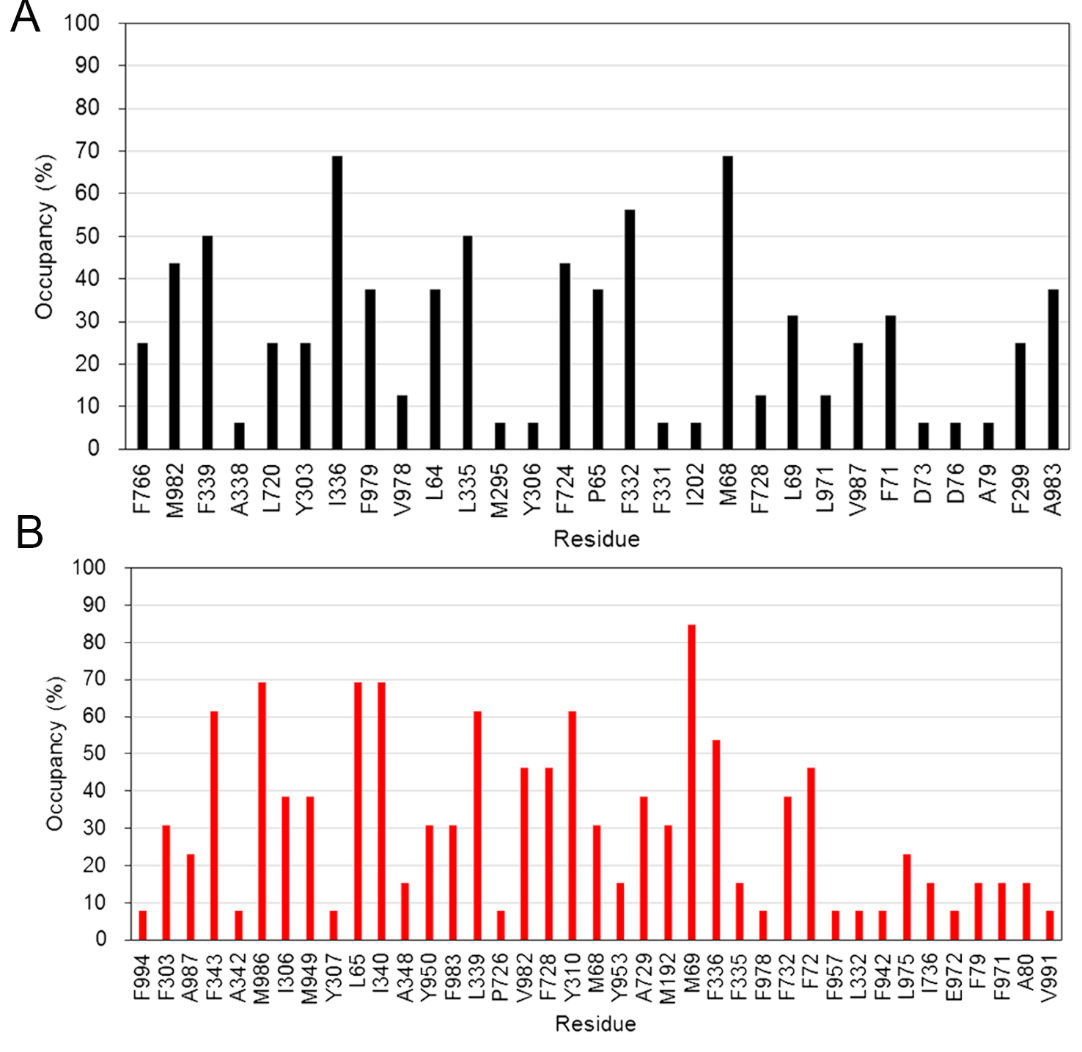

**Figure S7**. Residue occupancy for 30S transport in (A) mouse and (B) human P-gps.

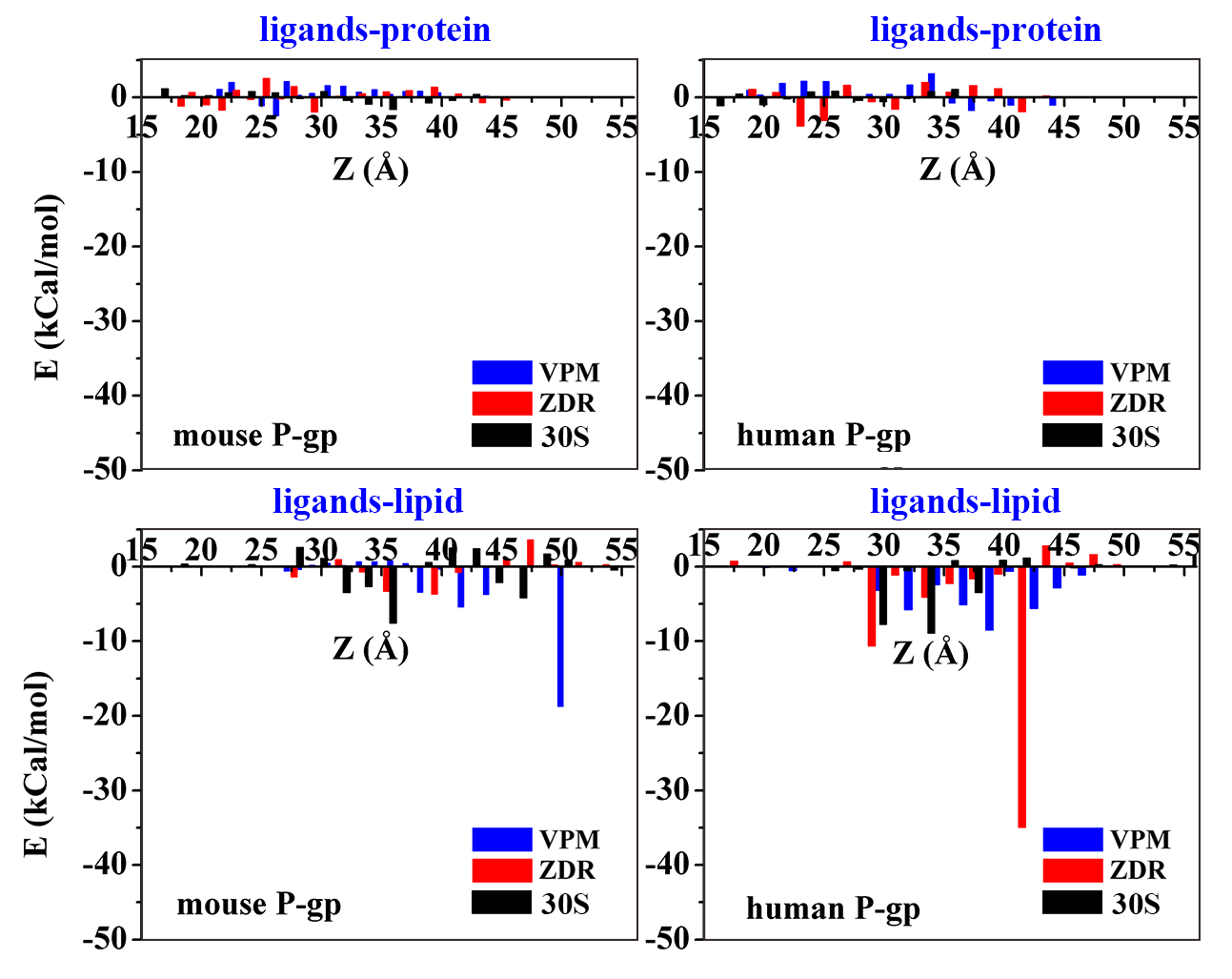

**Figure S8**. Electrostatic interaction energy between VPM/ZDR/30S and P-gp (top) or lipid (bottom) in mouse and human P-gp systems.

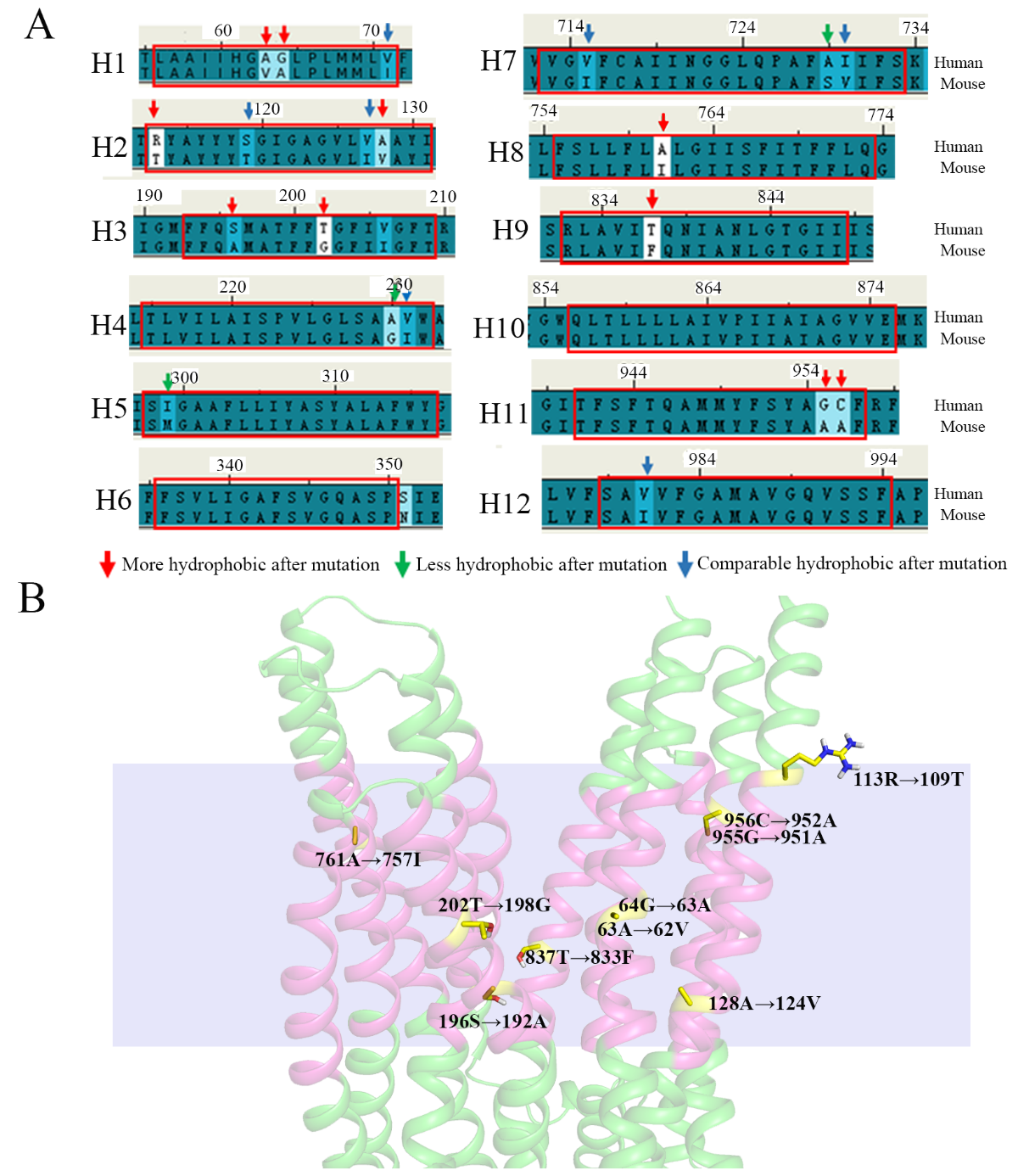

**Figure S9**. (A) Sequence alignment for transmembrane domain sequences of mouse and human P-gps. (B) The distribution of residues with higher hydrophobicity in mouse P-gp than in human P-gp in the transmembrane domain region.

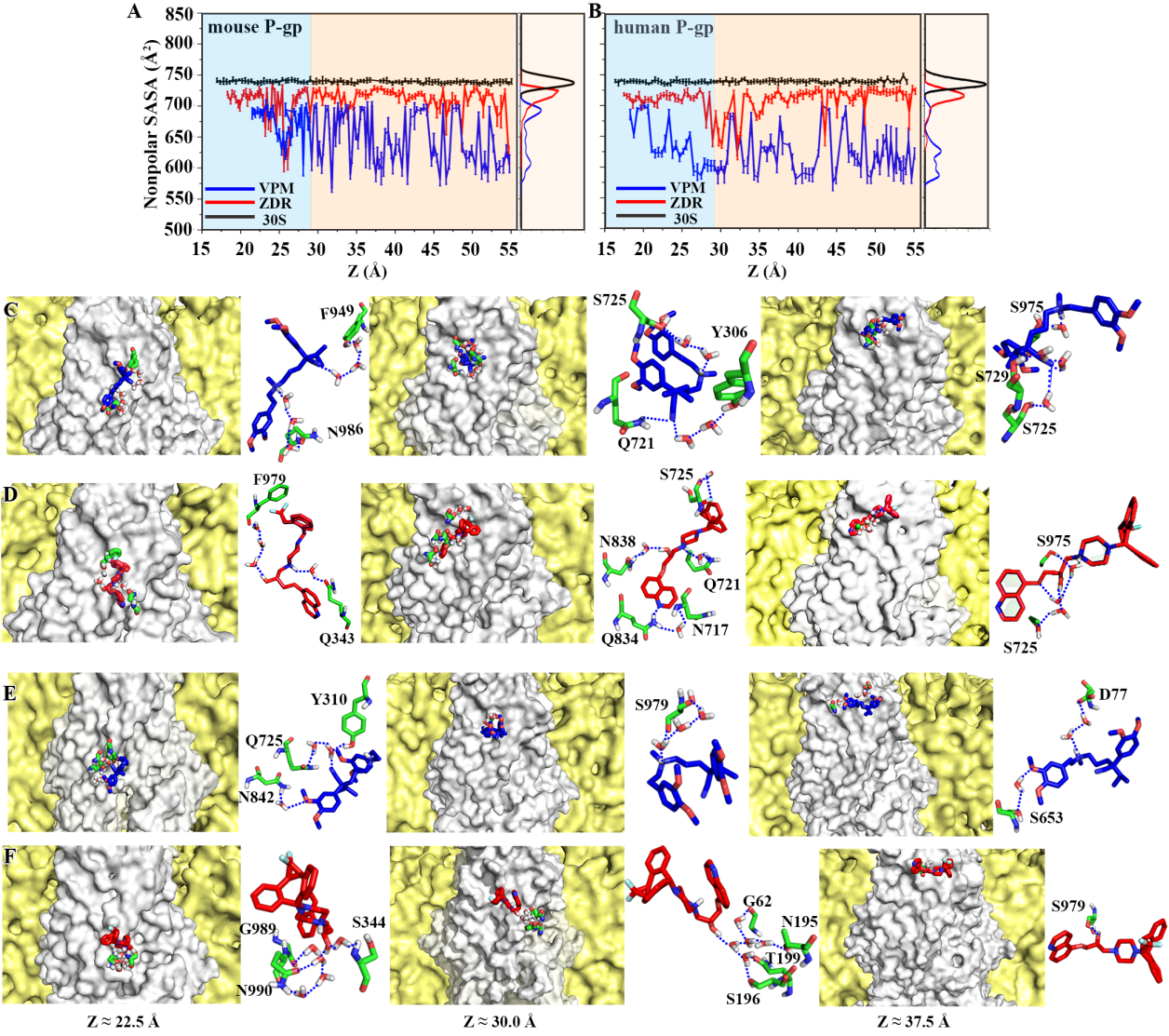

**Figure S10**. (A-B) Nonpolar solvent accessible surface area (SASA) of VPM/ZDR/30S along their transport pathways in mouse and human P-gps. (C-F) Representative snapshots from left to right indicating the presence of hydrogen bonding from P-gps towards the ligands for the systems of (C) VPM in mouse P-gp, (D) ZDR in mouse P-gp, (E) VPM in mouse P-gp, and (F) ZDR in human P-gp. The P-gps and surrounding lipid bilayer are represented by white and yellow surface. The hydrogen bonding is shown with dash lines and the ligands of VPM (blue) and ZDR (red), residues, and water molecules involved in hydrogen bonding are shown with stick representations.

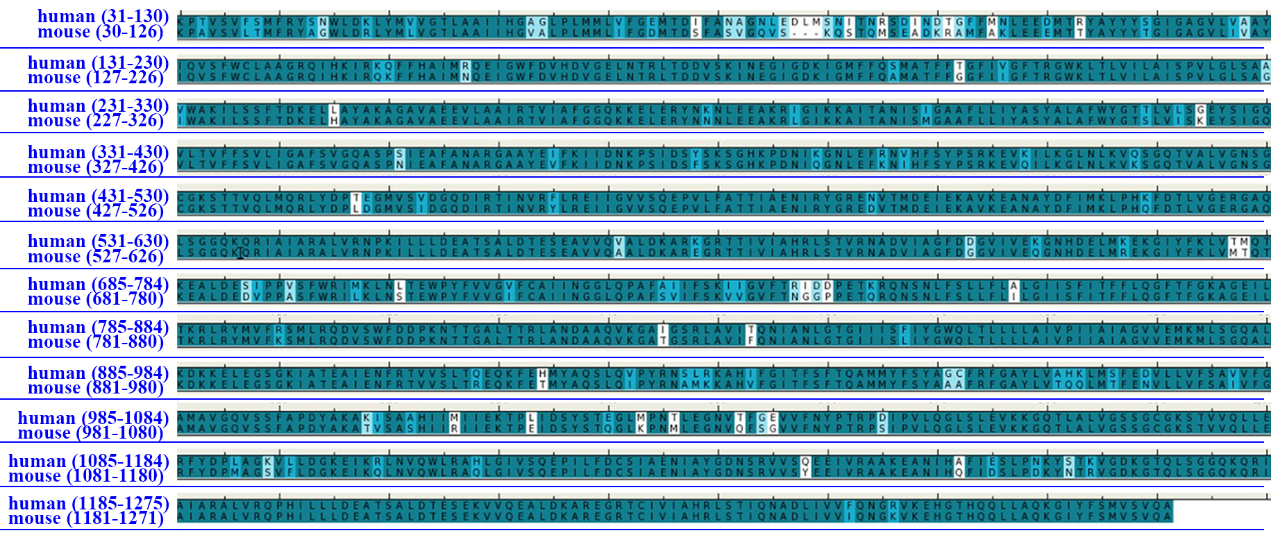

**Figure S11**. Sequence alignment of human and mouse P-gps.

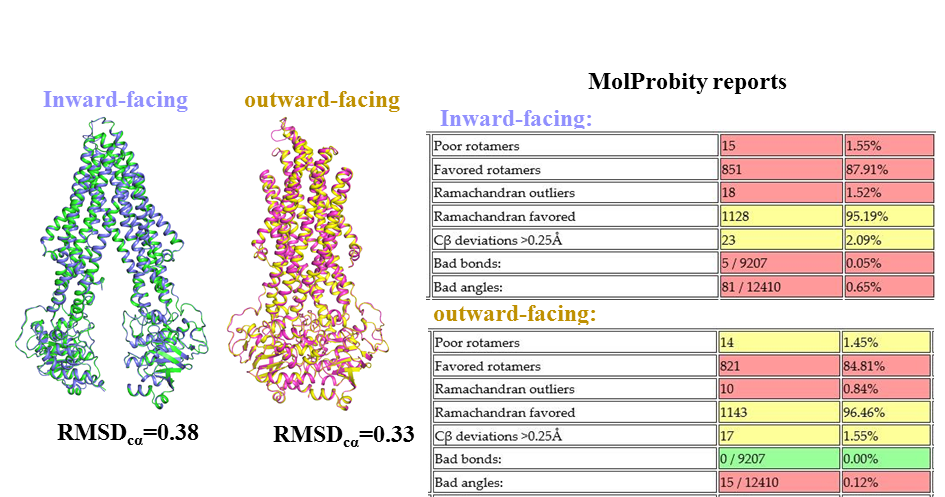

**Figure S12**. Homology modeling of inward-facing (purple) and outward-facing (yellow) conformations of human P-gp and the corresponding evaluation results by MolProbity.

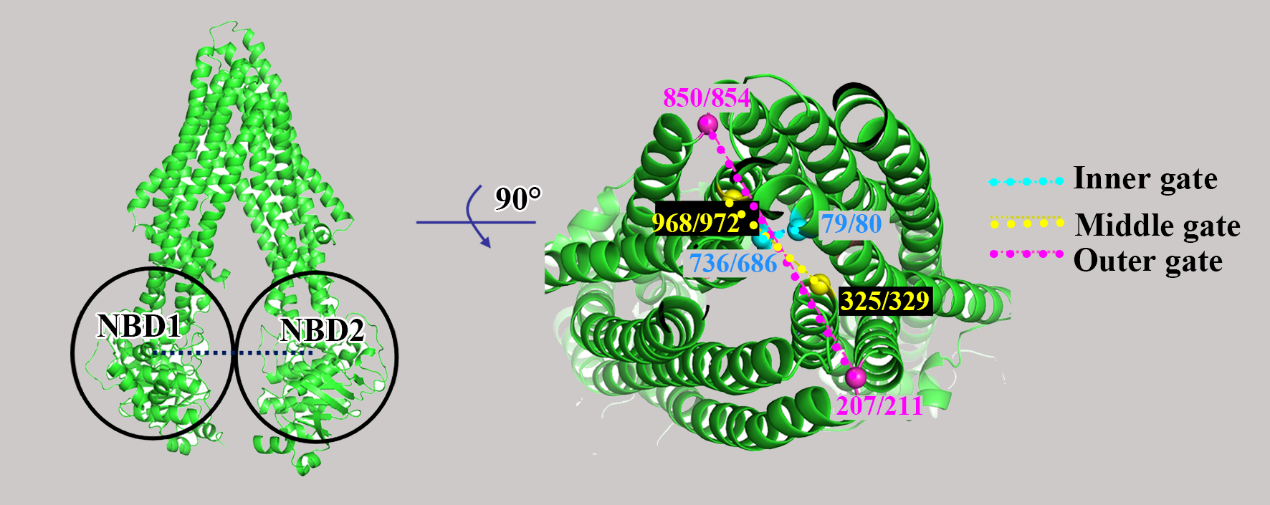

**Figure S13**. Schematic diagram indicating the reaction coordinates defining the conformational change of the nucleotide-binding domains (NBD1-NBD2), and the conformational change of the transmembrane domains including the inner gate (79/80-736/686), middle gate (325/329-968/972), and outer gate (207/211-850/854). The leading and trailing slash in brackets denote the residue index in mouse and human P-gps, respectively.

**Table S1.** Non-conserved residues in transmembrane domains of mouse and human P-gps.

| **Mouse** | 62V | 63A | 109T | 124V | 192A | 198G | 757I | 833F | 951A | 952A |
| --- | --- | --- | --- | --- | --- | --- | --- | --- | --- | --- |
| **Human** | 63A | 64G | 113R | 128A | 196S | 202T | 761A | 837T | 955G | 956C |
| **Mouse** | 226G | 295M | 725S | 70I | 115T | 123I | 227I | 711I | 726I | 977I |
| **Human** | 230A | 299I | 729A | 71V | 119S | 127V | 231V | 715V | 730I | 981V |

Residues with higher (lower) hydrophobicity in mouse P-gp than in human P-gp are in red (green) colors and those with similar hydrophobicity are in black.

**Table S2.** Detailed composition for P-gp-ligand complex systems under simulations.

|  | POPC | Water | Na+ | Cl- | Ligand | ATP | Mg^2+^ |
| --- | --- | --- | --- | --- | --- | --- | --- |
| Mouse P-gp | 397 | 50933 | 142 | 148 | VPM/ZDR/30S | 2 | 2 |
| Human P-gp | 388 | 58723 | 153 | 161 | VPM/ZDR/30S | 2 | 2 |

**Movie S1**. VPM transporting in mouse P-pg.

**Movie S1**. ZDR transporting in mouse P-pg.

**Movie S2**. 30S transporting in mouse P-pg.

**Movie S1**. VPM transporting in human P-pg.

**Movie S1**. ZDR transporting in human P-pg.

**Movie S1**. 30S transporting in human P-pg.
